# Fibroblast specialisation across microanatomy in a single-cell atlas of healthy human Achilles tendon

**DOI:** 10.1101/2025.10.17.683019

**Authors:** Carla J. Cohen, Jolet Y. Mimpen, Alina Kurjan, Claudia Paul, Shreeya Sharma, Lorenzo Ramos-Mucci, Chinemerem T. Ikwanusi, Ali Cenk Aksu, Tracy Boakye Serebour, Marina Nikolic, Kevin Rue-Albrecht, Christopher Gibbons, Duncan Whitwell, Tom Cosker, Steven Gwilym, Ather Siddiqi, Raja Bhaskara Rajasekaran, Harriet Branford-White, Adam P. Cribbs, Philippa A. Hulley, David Sims, Mathew J. Baldwin, Sarah J. B. Snelling

**Affiliations:** The Botnar Institute of Musculoskeletal Sciences, Nuffield Department of Orthopaedics Rheumatology and Musculoskeletal Sciences, University of Oxford, Oxford, UK; Centre for Computational Biology, MRC Weatherall Institute of Molecular Medicine, University of Oxford, Oxford, UK; Kennedy Institute of Rheumatology, Nuffield Department of Orthopaedics Rheumatology and Musculoskeletal Sciences, University of Oxford, Oxford, UK

**Author notes:** Equal contribution (Carla J. Cohen, Jolet Y. Mimpen). Equal contribution (Mathew J. Baldwin, Sarah J. B. Snelling).

## Abstract

Tendons are transitional tissues linking muscle to bone, enabling locomotion and fine motor control. The cellular biology across the Achilles tendon unit is poorly understood, yet critical for interpreting normal function and pathological changes across its microanatomically-defined functional zones. We generated a spatially-resolved transcriptomic atlas of human Achilles tendon, sampling the tendon-bone junction (enthesis), midbody, myotendinous junction, and adjoining muscle. Six fibroblast subtypes were identified, with distinct transcriptional profiles and spatial distributions, suggesting specialised functional roles across the tendon-muscle unit. Two dominant fibroblast types were specifically positioned in the tendon mid-substance and paratenon (vessel-rich region surrounding the tendon fibrils); other populations included perineural, myotendinous junction-specific, muscle-specific, and lining-layer fibroblasts. These findings demonstrate how cellular diversity across a transitional tissue may underlie microanatomical-specific roles. This atlas provides a foundation for understanding cellular functions across the tendon-muscle unit and will be essential for comparisons with diseased tissue, identifying pathogenic mediators and treatment targets for autoimmune and degenerative pathologies of the Achilles tendon.

## INTRODUCTION

The Achilles tendon is the largest the human body. It is an energy-storing tendon and is crucial for mobility, transferring mechanical forces through linking the calcaneal bone of the ankle to the Gastrocnemius and soleus muscles of the lower leg (1). Tendon, muscle, and bone are classically regarded as distinct tissue types with unique cell types and functions. However, tendons transition through microanatomical zones with specific biological and mechanical requirements, particularly at bone (enthesis) and muscle (myotendinous) junctions (Figure 1). At a cellular level, these microanatomies are likely to require different resident cell types and relative cellular compositions to maintain tendon health in response to normal loading and to orchestrate the response to injury and disease (2). Achilles tendon ruptures are the most common tendon injury of the lower extremities, and predominantly occur in the midbody of the tendon but also at the myotendinous junction (MTJ) and enthesis (3, 4). The Achilles can also be affected by enthesopathies, inflammation of the enthesis, including ankylosing spondylitis and psoriatic arthritis (5). A deeper understanding of the transitional tendon microanatomy in health, and in particular the specific cellular composition of each zone, is needed to understand its contribution to normal mechanical processes and in disease (6).

**Figure 1:**
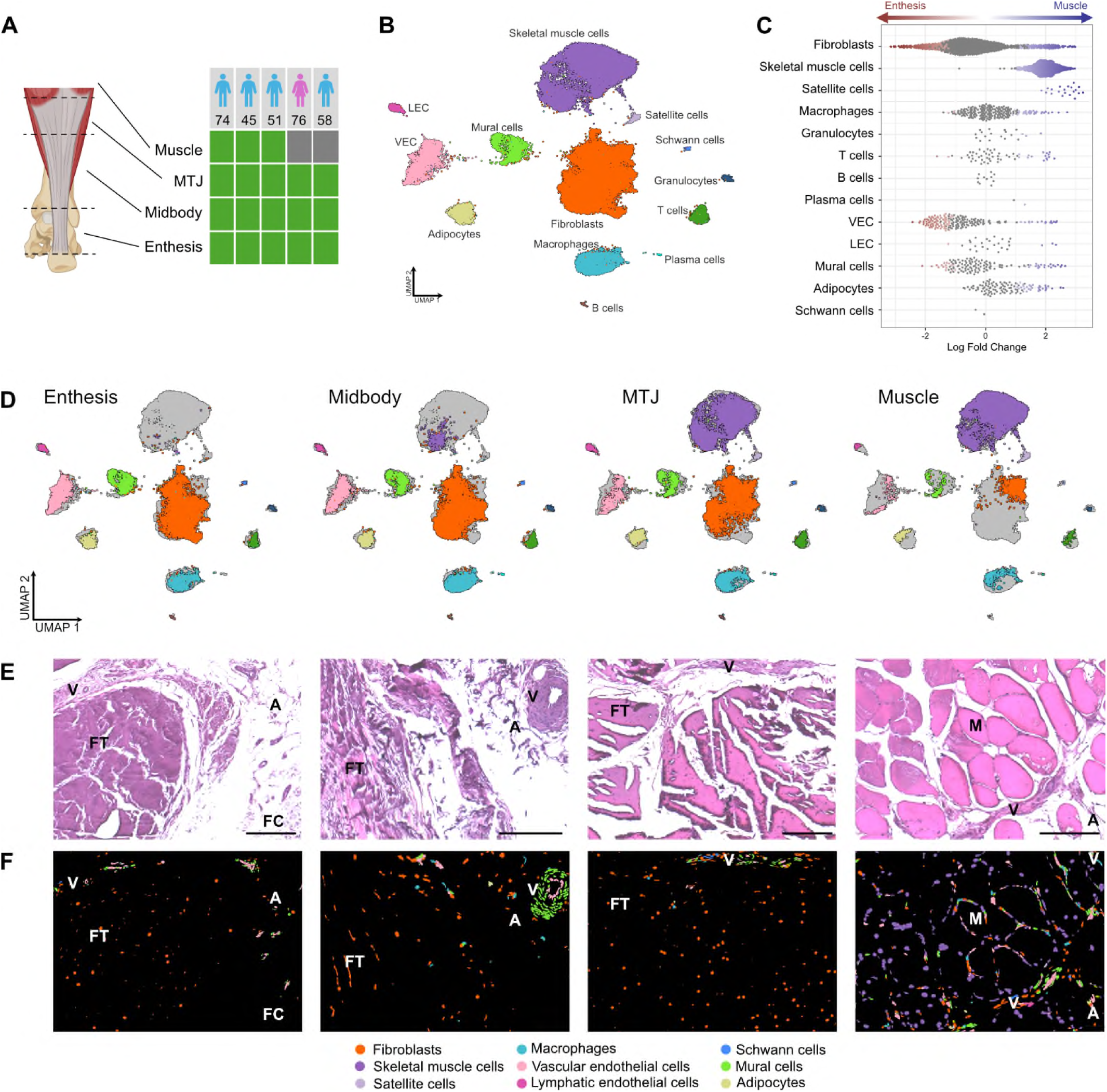
Cell composition varies considerably across Achilles tendon microanatomy. **(A)** Schematic of the lower leg and human Achilles tendon showing tissue regions sectioned for single-nucleus RNA-seq, spatial transcriptomics and imaging at three Achilles tendon microanatomies (enthesis, midbody, MTJ) and adjoining muscle. Table shows sample inclusion, sex and age of participants (n=5). (**B**) UMAP representation of broad cell types found in Achilles tendon and adjoining muscle using snRNA-seq. (**C**) Beeswarm plot showing differential abundance of each broad cell type across microanatomy calculated using MiloR. Each dot represents one cellular neighbourhood comprising tens-hundreds of similar cells; red and blue colour indicates neighbourhoods with significantly greater abundance at the enthesis or muscle respectively. (**D**) UMAP representation of broad cell types showing cells present at each microanatomy (same colours as B). (**E**) Representative spatial transcriptomics plots overlaid onto post-Xenium H&E images from each microanatomy. Labels: FT – Fascicular tendon; V – vessel; A – adipose tissue; FC – fibrocartilage; M – muscle. Broad cell annotations were predicted from the snRNA-seq data; colours as in B. Scale bar 250 µm.

Histologically, the normal midbody tendon consists of collagen I-rich fibrils, that tightly pack to form parallel bundles or fascicles (2, 7). The major cell type within this region is typically regarded to be “tenocytes” – fibroblasts that produce and turnover the extracellular matrix that comprises the bulk of tendon tissue. Collagen fascicles are surrounded by an endotenon which acts as a conduit for nerve, blood and lymphatic vessels, and is an extension of the epitenon and paratenon (together called peritendon) that surrounds the tendon body (1, 8). Towards the MTJ, skeletal muscle fibres and collagen I fibrils are seen to interdigitate, maximising surface area and ensuring strong connections between the tendon and muscle tissue (9). Within the enthesis region, collagen fibre alignment decreases, collagen II and proteoglycan levels increase, as does mineralisation status, with chondrocyte-like cells also present (10). These transitions across tendon microanatomy enables effective force transfer between muscle and bone.

Single-cell and spatial transcriptomic studies have identified functional tissue niches comprising specific cellular compartments, for example in the gut and kidney (11, 12). The complexity of tendon cellular composition has been revealed by recent single-cell studies of healthy human hamstring, quadriceps and supraspinatus, and developmental tendons (13–22). While fibroblasts remain the predominant cell type in these healthy human tendons, distinct fibroblast subtypes were also identified, alongside endothelial cells, immune cells, adipocytes and nervous system cells. Comparison with diseased tissue highlighted alterations in cell activation states in tendon tears. Single-cell transcriptomic analysis of healthy Achilles tendon in 6-week-old mice also described fibroblast, endothelial, and immune populations (23). However, a detailed analysis of the healthy human Achilles tendon, with its unique energy-storing properties, and the relation of cells to their spatial position within tendon microanatomy remains absent.

Here we present the first transcriptomic and spatial atlas of healthy human Achilles tendon which identifies specific cell types across the tendon-muscle unit, comparing the enthesis, midbody and MTJ of the tendon and adjacent muscle region. We show that the composition, transcriptional profile, and tissue location of fibroblasts vary considerably across Achilles tendon microanatomy; these differences seem to be associated with six specialist fibroblast subtypes, each with unique ligand-receptor interaction profiles and functions. This comprehensive description of cell composition in healthy human Achilles tendon homeostasis will be an invaluable reference for future comparative studies of the cellular pathogenesis of Achilles tendinopathy and in regenerative medicine approaches targeting the Achilles tendon. The new knowledge of fibroblast composition, which changes with microanatomical zone, will spatially inform mechanistic understanding and therapeutic targeting of tendon health and pathology.

## RESULTS

### Cellular composition of the Achilles tendon-muscle unit reflects its **microanatomical structure**

To study the tissue morphology and cell types present in the Achilles tendon-muscle unit we collected tissue from microanatomical sites: tendon proximal to the enthesis, tendon midbody, tendon proximal to the MTJ, and adjoining muscle (Figure 1A). We used histological staining to compare tissue features across microanatomy (Figure S1). Across the whole tendon, densely packed fascicular tendon was visible, interspersed with inter-fascicular matrix. Surrounding this was peritendon rich in blood vessels, nerves and adipocytes. At the enthesis the junction between tendon fibrils and acellular fibrocartilage was clearly visible. The muscle-tendon interaction was apparent at the MTJ, although muscle fibres extended into the tendon midbody. The muscle tissue section showed skeletal muscle fibres alongside a stromal section encompassing vessels and nerves, much like the peritendon.

We isolated nuclei from tissue across the Achilles-tendon unit from five patients (Figure 1A, Table 1) and performed single nucleus RNA sequencing (snRNA-seq). After quality control and integration (Figure S2), we obtained 64,862 nuclei and assigned each to its putative cell type of origin. Using canonical markers we found 13 broad cell types including fibroblasts (*DCN*, *COL1A2*), skeletal muscle cells (*TTN*, *NEB*) and their precursor satellite cells (*PAX7*, *CALCR*), immune cells comprising macrophages (*CD163*, *MRC1*), T cells (*THEMIS*, *CD247*), granulocytes (*KIT*, *IL18R1*), B cells (*BLNK*, *FCRL1*)and plasma cells (*JCHAIN*, *XBP1*), and non-fibroblast stromal cells comprising vascular endothelial cells (VEC; *PECAM1*, *VWF*), lymphatic endothelial cells (LEC; *PROX1*, *PKHD1L1*), mural cells (*NOTCH3*, *MYO1B*) and adipocytes (*ADIPOQ*, *AQP7;* Figure 1B, Figure S3). We sought to determine the variation in cell type abundance across the tendon-muscle unit using quantitative differential abundance testing (Figure 1C) and analysis of cell proportions (Figure 1D, Figure S4). Spatial transcriptomics by 10X Xenium with post-analysis H&E staining was performed to facilitate visualisation of the cell positioning within the densely fibrous tendon tissue and muscle (Figure 1E-F). Fibroblasts were the most numerous cell type (37% of nuclei) and were present throughout the length of the tendon from the junction with fibrocartilage to the MTJ, and in muscle. Spatial transcriptomics confirmed that fibroblasts (red) were located in the peritendon and fascicular tendon, as well as adjacent to muscle fibres (Figure 1F). From the differential abundance testing, it appeared that fibroblasts had a wide spread of neighbourhood enrichment, suggesting that there might be fibroblast subsets specific to each microanatomical zone (Figure 1C). Unsurprisingly, skeletal muscle cells and satellite cells were highly enriched at the muscle and MTJ and were almost absent from the enthesis-proximal tendon (Figure 1C), and in spatial data the myonuclei could be seen surrounding muscle fibres (Figure 1F). The peritendon structure and cellular composition seemed consistent across microanatomy and contained adipocytes (yellow), mural cells (green) and VEC (pink) cells visible around blood vessels, Schwann cells (dark blue) in peripheral nerve bundles and small numbers of immune cells (mostly macrophages, light blue) (Figure 1EF). The muscle samples also contained cells associated with blood vessels, nerves and lymphatic structures. Small populations of T cells, B cells, plasma cells and granulocytes were identified in the snRNA-seq data but rarely detected spatially. Overall, these findings confirmed the cellular composition and spatial position of broad cell types, and led us to consider whether smaller cell subtypes, particularly fibroblasts, might have specialist locations across the Achilles tendon unit.

**Table 1.** Summary of patient demographics, including de-identified patient ID, sex, age (in years, affected side, and whether muscle was collected.

| Patient ID | Sex | Age (yrs) | Affected side | Muscle collected |
| --- | --- | --- | --- | --- |
| MSK0785 | Male | 74 | Left | Yes |
| MSK1250 | Male | 45 | Left | Yes |
| MSK1556 | Male | 51 | Right | Yes |
| MSK1687 | Female | 76 | Left | No |
| MSK1691 | Male | 58 | Right | No |

### Distinct fibroblast subsets with unique functional properties are found **across the Achilles tendon-muscle unit**

Next, we identified which genes were differentially expressed (DE) across the Achilles tendon microanatomy within each broad cell type. While most cell types had relatively few DE genes (<1000), fibroblasts had over 5000 DE genes (Figure S5A). By clustering the genes according to their expression profiles across microanatomy, we found ten distinct groups of DE genes from fibroblasts (Figure 2A). The two largest clusters were those with the highest expression in the muscle (cluster 1; 761 genes) or at the enthesis (cluster 2; 491 genes). We also identified tendon-specific gene clusters with high expression across tendon and low expression in muscle (clusters 4, 6 and 8; 245 genes), and muscle-specific gene clusters with high expression in muscle and low expression in tendon (clusters 5, 9 and 10; 133 genes). Pathway analysis on these combined gene clusters revealed enrichment of distinct biological pathways for each gene expression pattern, such as muscle-related pathways for gene clusters with high expression in muscle, and adhesion-related pathways for clusters with high expression in enthesis (Figure 2B). These findings led us to consider whether annotation at finer resolution could identify fibroblasts with more specialist locations and functions.

**Figure 2:**
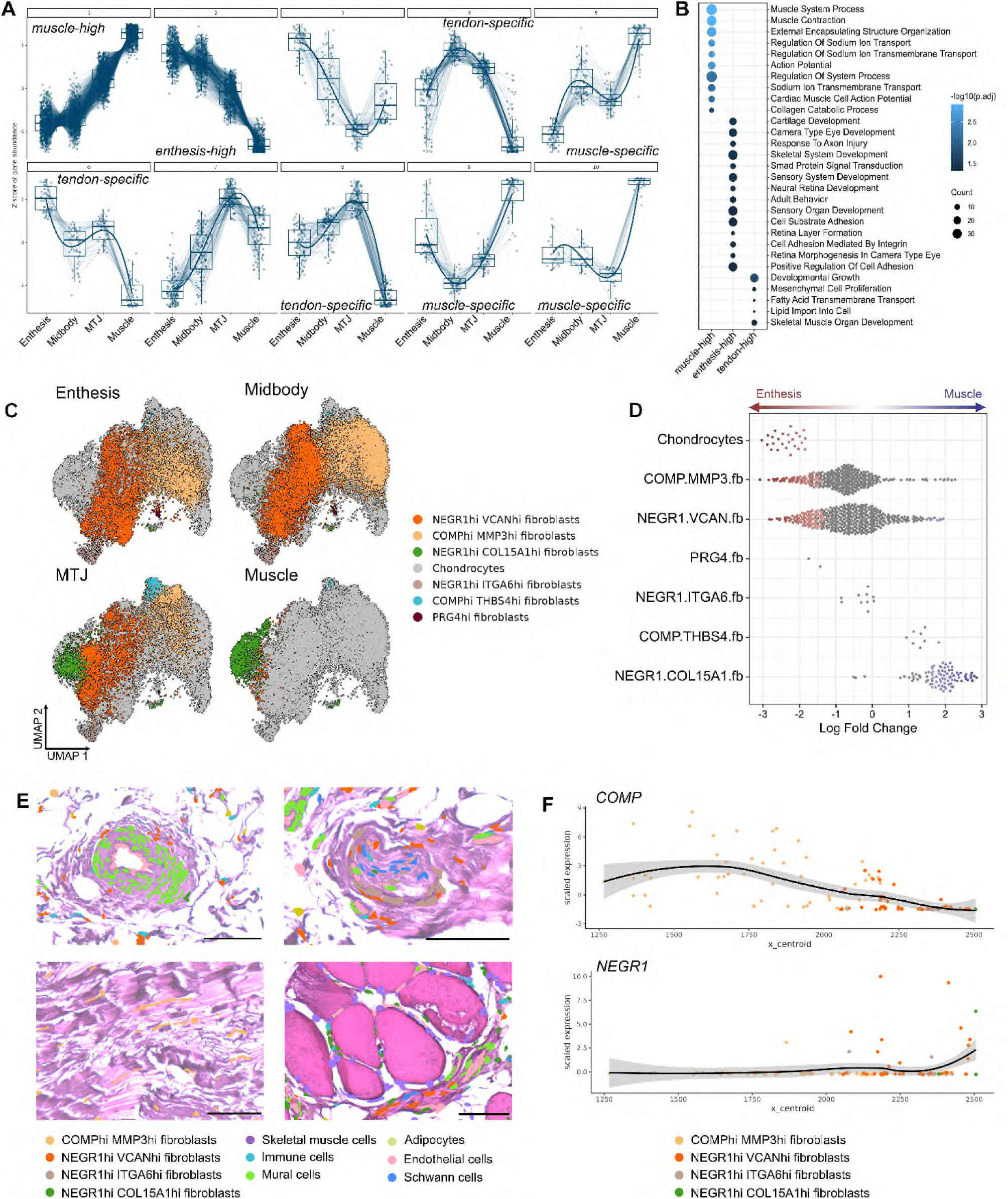
Phenotypic and functional characterisation of fibroblast subtypes. **(A)** Scaled gene expression in fibroblasts clustered by expression patterns across tendon microanatomy. Groups of genes with particular cluster profiles are indicated (muscle-high, enthesis-high, tendon-specific and muscle-specific). (**B**) Dotplot showing GO:BP pathways enriched in clusters with muscle-high expression, enthesis-high expression or tendon-high expression. Although 133 muscle-specific genes were found, these were not enriched in any pathways. (**C**) UMAP representation of the fibroblast subset showing fine cell type annotations split by microanatomy. (**D**) Beeswarm plot showing differential abundance of each fibroblast subtype across microanatomy calculated using MiloR. Each dot represents one cellular neighbourhood comprising tens-hundreds of similar cells; red and blue colour indicates neighbourhoods with significantly greater abundance at the enthesis or muscle respectively. (**E**) Spatial transcriptomics plots showing fibroblast subtypes overlaid onto post-Xenium H&E images. Upper left: NEGR1hi VCANhi fibroblasts positioned near a blood vessel in peritendon from tendon midbody; Lower left: COMPhi MMP3hi fibroblasts in the fascicular tendon from midbody; Upper right: NEGR1hi ITGA6hi fibroblasts surrounding a peripheral nerve bundle in peritendon from tendon midbody; Lower right: NEGR1hi COL15A1hi fibroblasts interspersed with muscle fibres and vessels in muscle. (**F**) Dotplot showing scaled expression of *COMP* and *NEGR1* according to fibroblast subtype along a cross-section of tendon midbody (x-centroid). Black line shows smoothed expression calculated by ggplot2 geom_smooth() with standard error in grey.

Fine-resolution annotation of fibroblasts in the snRNA-seq data identified six fibroblast subtypes and chondrocytes (Figure 2C, Figure S5B-H). The five biggest clusters could be broadly classed into two groups, expressing *NEGR1* and *COMP*, and more specifically using additional markers *VCAN*, *MMP3*, *COL15A1, ITGA6*, and *THBS4.* In addition, two smaller clusters were found and identified as chondrocytes (*ACAN*) and PRG4+ fibroblasts (*PRG4*). Each subtype expressed genes enriched in specific biological pathways (Figure S5I). Differential abundance analysis of the fibroblast subsets which, along with variation in relative proportion of nuclei, suggested striking differences in the distribution of fibroblast subtypes across microanatomy (Figure 2C, D, Figure S5J-L). Expression of highly DE genes across microanatomy seemed to correlate with genes that are highly expressed in fibroblast subtypes that are mostly present in that microanatomy (Figure S5M), suggesting that the variation is due to alterations in cell composition rather than transcriptional changes across microanatomy within a particular fibroblast.

To uncover the spatial location and organisation of these specialist fibroblasts within the tendon and muscle tissue, we used spatial transcriptomics (Figure 2E). The most abundant fibroblast populations were NEGR1hi VCANhi fibroblasts (44% of nuclei) and COMPhi MMP3hi fibroblasts (40% of nuclei), which were both present in all tendon microanatomies but absent from muscle. NEGR1hi VCANhi fibroblasts were present specifically in the peritendon adjacent to blood vessels and nerves while COMPhi MMP3hi fibroblasts were found in the fascicular/inter-fascicular tendon (Figure 2E). Consistent with this, COMPhi MMP3hi fibroblasts expressed genes in pathways related to extracellular matrix adhesion (Figure S5H). Expression of the key marker genes *COMP* and *NEGR1*correlated spatially with the location of COMPhi MMP3hi and NEGR1hi VCANhi fibroblasts across a cross-section of the field of view in midbody and confirmed that these two populations have non-overlapping spatial locations in the tendon tissue (Figure 2F). NEGR1hi ITGA6hi fibroblasts were found surrounding peripheral nerves in the tendon and muscle, so appear to be perineural fibroblasts (Figure 2E). NEGR1hi COL15A1hi fibroblasts were positioned surrounding the muscle fibres in the muscle tissue section (Figure 2E). The small number of COMPhi THBS4hi fibroblasts were predominant at the MTJ in snRNA-seq, but not detected in the spatial data (Figure 2C-D, Figure SI). Finally, chondrocytes (*ACAN*+ cartilage matrix-producing cells) and PRG4hi fibroblasts were detected in the snRNA-seq data but not in the spatial data, with chondrocytes specific to the enthesis and PRG4hi fibroblasts present in the tendon but not muscle (Figure 2C-D, Figure S5K-L).

We performed SCENIC analysis (24) and identified transcriptional networks (regulons) specifically enriched in each fibroblast subset providing further evidence of their functional specialisation (Figure S5N). For example, the IRF3 regulon, known to regulate collagen production in fibrosis (25), was enriched in COMPhi MMP3hi fibroblasts, whereas the MYF6 regulon was enriched in MTJ/muscle-resident COMPhi THBS4hi and NEGR1hi COL15A1hi fibroblasts, perhaps conferring contractile properties in these cells (26).

### Fine annotation of non-fibroblast cell types

Fine annotation of the non-fibroblast cell types - muscle cells, immune cells and non-fibroblast stromal cells – revealed 22 distinct subtypes in snRNA-seq data (Figure 3, Figure S6-8). Within the non-fibroblast stromal cells, the adipocytes, LEC, and Schwann cells did not display any further subsets, but arteriolar (*PODXL, EFNB2, NEBL*) and venular VEC (*NOSTRIN, MYRIP, GNA14*) were distinguishable as well as pericytes (*PDGFRB, EGFLAM, RIMS1*) and vascular smooth muscle cells (vSMC; *MYH11, RCAN2, LMOD1*) (Figure 3A, Figure S6). Differential abundance analysis suggested moderate changes in the distribution of cell neighbourhoods across the Achilles tendon-muscle unit, especially pericytes which were highly enriched towards muscle (Figure 3A). All stromal cells were observed in H&E-defined cross-sections of veins, arteries, nerves, lymphatic structures and adipose in tendon and muscle using spatial transcriptomics (Figure 3D-G).

**Figure 3:**
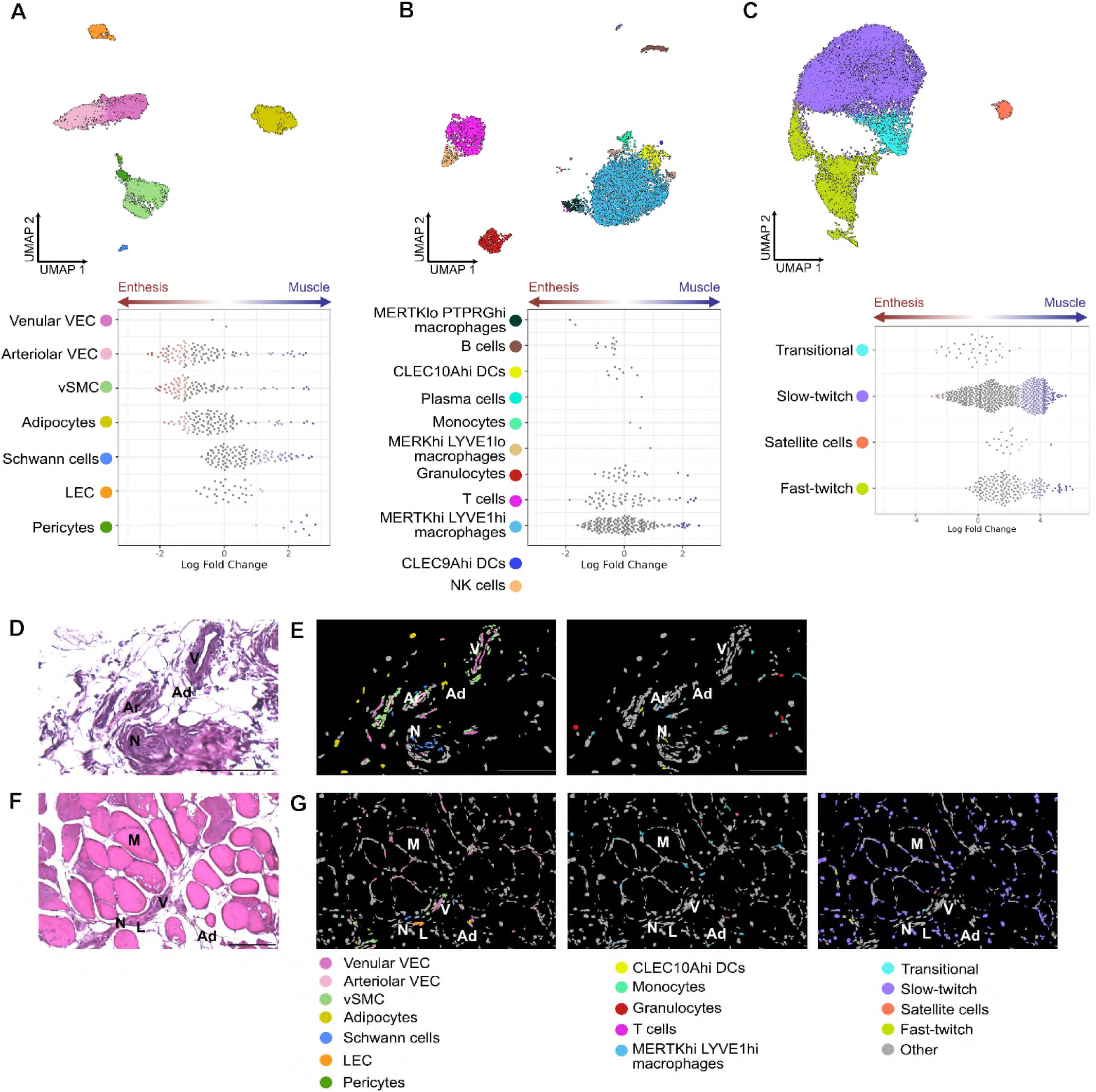
Fine annotation of non-fibroblast cell types identified in Achilles tendon. (**A-C**) UMAP representations (upper) and Beeswarm plots (lower) from (**A**) stromal subset (**B**) immune subset and (**C**) muscle subset. UMAP representations are coloured by fine annotation level; Beeswarm plots show differential abundance of each finely annotated cell type across microanatomy calculated using MiloR. Immune cell subtypes CLEC9Ahi DCs and NK cells did not form distinct neighbourhoods since they were few in number. (**D-G**) H&E staining and spatial transcriptomics images showing tissue position of finely annotated cell types. (**D**) H&E image from Achilles tendon midbody, showing blood vessels, adipocytes and a peripheral nerve bundle in the peritendon. (E) Spatial transcriptomics plots of the same field of view as (D) coloured by stromal (left) or immune (right) cell types. (**F**) H&E image from muscle, showing muscle fibres, veins, nerves and lymphatic vessels. (**G**) Spatial transcriptomics of the same field of view as (F) coloured by stromal (left), immune (middle) or muscle (right) cell types. Other cell types are coloured grey, and scale bars are 200 µm. Labels: V – vein; Ar – artery; N – nerve bundle; Ad – adipose; M – muscle; L – lymphatic vessel.

Macrophages were the most abundant immune cells in snRNA-seq data, comprising a major population of tissue-resident MERTKhi LYVEhi macrophages alongside MERTKhi LYVE1lo and MERTKlo PTPRGhi macrophage populations (27). Fine annotation also revealed monocytes, CLEC10Ahi and CLEC9Ahi dendritic cells (DCs) and NK cells, alongside T cells, B cells, plasma cells and granulocytes (Figure 3B, Figure S7). While some differential abundance analysis suggests MERTKlo PTPRGhi macrophages were enriched towards the enthesis and some small neighbourhoods of granulocytes, T cells, and MERTKhi LYVE1hi macrophages were enriched towards the muscle, most immune cell types did not show striking changes across microanatomies in snRNA-seq data (Figure 3B). The major immune cell groups (DC, monocytes, Granulocytes, T cells and macrophages) were detectable in the spatial transcriptomics data, in or adjacent to blood vessels in both tendon and muscle (Figure 3D-G).

The muscle cell populations predominantly comprised classical slow-twitch (type I) and fast-twitch (type II) myocytes. Transitional myocytes were annotated by expression of both slow and fast twitch markers as well as *COL22A1*, an MTJ marker (28), and satellite cells (myocyte precursors) were identified using *PAX9* (Figure 3C, Figure S8). Differential abundance analysis revealed that neighbourhoods of each subset were enriched towards the MTJ and muscle (Figure 3C). Slow and fast-twitch muscle nuclei were observed spatially surrounding distinct muscle fibres (Figure 3G), and small numbers of transitional muscle cells and satellite cells could also be seen in the muscle section. A bundle of slow-twitch muscle fibres was seen in the tendon midbody section, indicating that the muscle fibres can extend deeply into the tendon structure (Figure S8E). Unexpectedly, unlike in the histological staining, we did not detect any muscle cell types in the MTJ spatial transcriptomics section, which is likely a sampling issue.

### Cell-cell interactions are dependent on cell composition and fibroblast **function**

Having identified differences in spatial positioning of fibroblasts across tendon microanatomy, we next identified predicted cell-cell interactions between all the cell types found in the Achilles tendon-muscle unit (Figure 4A). Overall, there were more ligand-receptor interactions in muscle than tendon, and fibroblasts were the cell type with the most interactions. The number of interactions appeared to relate to cell composition, with more abundant cells having more interactions at a particular microanatomy. For example, chondrocytes showed strong interactions with NEGR1hi VCANhi fibroblasts only in the enthesis, whereas NEGR1hi COL15A1hi had more interactions with COMPhi THBS4hi fibroblasts in MTJ and muscle. We therefore focussed on interactions between specific fibroblast subtypes and non-fibroblast cells (Figure 4B-D). Very few interactions were seen between fibroblasts and immune cells or muscle cells in this healthy tissue (Figures S9). NEGR1hi COL15A1hi fibroblasts (resident in muscle), and NEGR1hi VCANhi fibroblasts (in vessel-rich peritendon) express *SCARA5* (scavenger receptor 5 class A), which is predicted to interact with *VWF* (Von-Willebrand Factor) on vascular endothelial cells (Figure 4B). In addition, the NEGR1hi COL15A1hi fibroblasts express several ECM genes (*MMP2*, *COL4A1*, *COL4A2*) which are predicted to bind receptors on the vascular endothelial cells (*PECAM1*, *CD93*) (Figure 4C). NEGR1hi ITGA6hi fibroblasts, found in nerve bundles, showed specific interaction with Schwann cells in the nerve via *NRXN1* (Neurexin-1) – *NLGN1* (Neuroligin-1) (Figure 4D). This analysis reveals that the interactions between fibroblasts and other cells are influenced by their spatial position in the tissue, providing additional insights into their putative functions.

**Figure 4:**
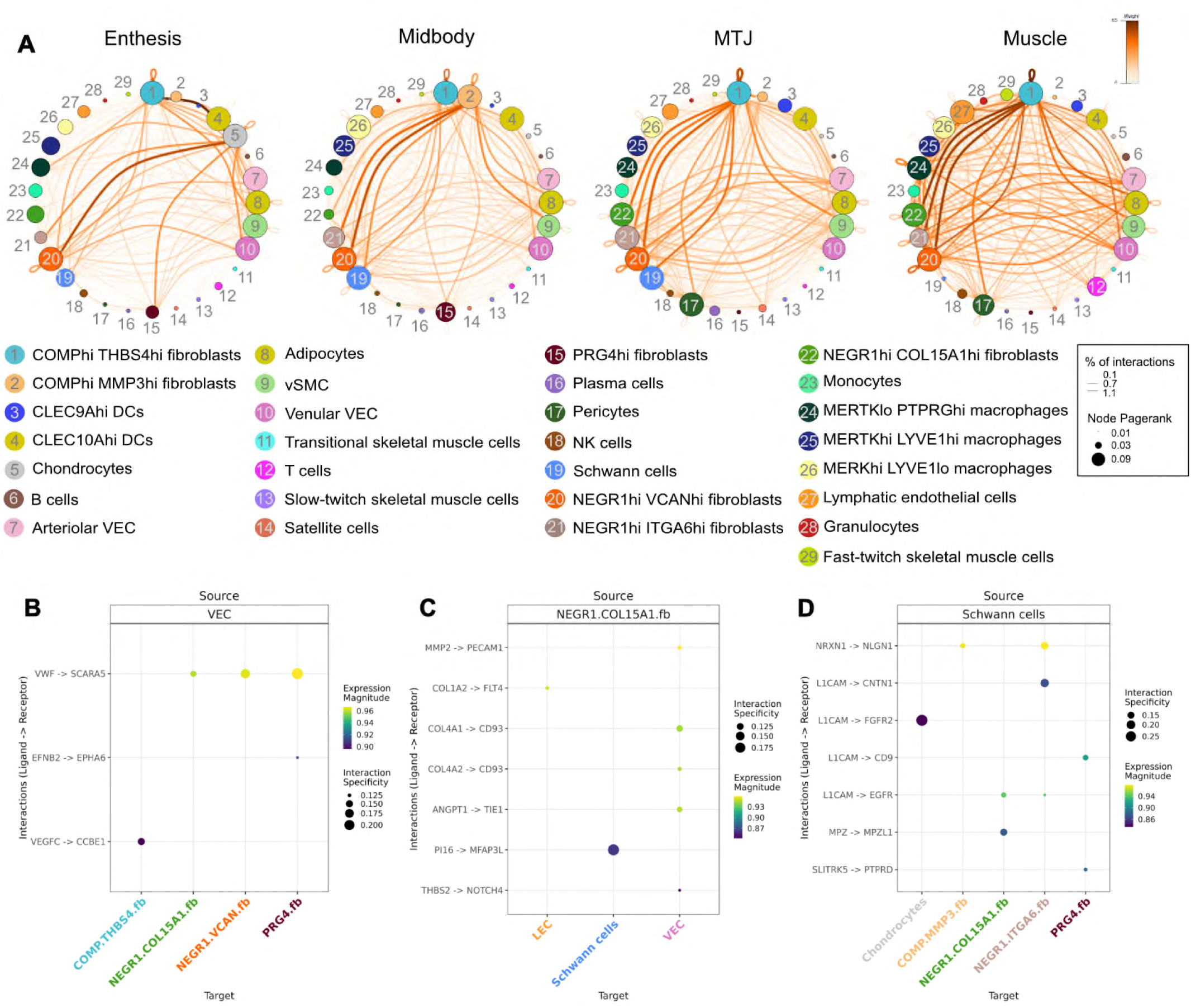
Predicted cell-cell interactions across cell types in healthy Achilles tendon. **(A)** Cell-cell interaction plots showing ligand-receptor interactions at each microanatomy calculated using LIANA. Each node on the circumference represents a cell type with the size of the dot related to the number of interactions with that cell type (Pagerank). Interactions are depicted by lines between the dots with the thickness representing the percent of total interactions and colour representing the weight of the interaction. Dotplot show specific ligand-receptor interactions between (**B**) VEC and fibroblasts, (**C**) NEGR1hi COL15A1hi fibroblasts and stromal cells, (**D**) Schwann cells and fibroblasts. Abbreviations: COMP.THBS4.fb – COMPhi THBS4hi fibroblasts; NEGR1.ITGA6.fb – NEGR1hi ITGA6hi fibroblasts; NEGR1.COL15A1.fb – NEGR1hi COL15A1 fibroblasts; COMP.MMP3.fb – COMPhi MMP3hi fibroblasts; NEGR1.VCAN.fib - NEGR1hi VCANhi fibroblasts.

## DISCUSSION

This study presents the first cellular atlas of the healthy human Achilles tendon-muscle unit in which macroscopic differences in tissue architecture were mirrored at the cellular level, reflecting the functional properties of each microanatomical zone. Notably, six distinct fibroblast subtypes were defined, each with specific transcriptional profiles, alongside stromal, immune and muscle cell phenotypes, consistent with those reported in other tendon studies (13–18, 20, 28). By including high-resolution spatial transcriptomic data, we have been able to identify the exact locations of these fibroblast subsets within the tendon and muscle tissue relative to the tendon matrix, vessels, nerves, muscle fibres, and non-fibroblast cell types. Some fibroblast subsets are enriched within a particular microanatomical zone and have unique cell-cell interactions. Taken together, we were able to predict functions of each fibroblast subtype that contribute to the microanatomical niches across the tendon-muscle unit.

At the expression level, five of the fibroblast subsets we identified within the Achilles tendon fell into two broad groups defined by the expression of *COMP* and *NEGR1* (Figure S5F-G); these two subsets have similar gene expression profiles to the MKX+ (*COMP*, *ITGA10*, *THBS4, COL12A1*) and PDGFRA+ (*NEGR1*, *FBN1*, *DCLK1*) fibroblasts found in hamstring tendon, respectively (14). COMPhi MMP3hi fibroblasts (40% of fibroblasts) and NEGR1hi VCANhi fibroblasts (44% of fibroblasts) were present across all tendon microanatomies but absent from the muscle, so are likely to be “tenocytes” or tendon-specific fibroblasts. COMPhi MMP3hi fibroblasts were located in the fascicular tendon, expressed genes related to matrix adhesion (Figure 2) and were enriched for regulons *FLI1* and *IRF3* (Figure S5), which are known to regulate collagen production (25, 29). As such, these fibroblasts appear to be required for maintenance of the ECM that comprises the fascicular tendon and are similar to FBLN1hi fibroblasts (*COMP*, *FBLN1*, *CILP*) in healthy quadriceps tendon (13). NEGR1hi VCANhi fibroblasts expressed genes related to cell morphogenesis consistent with their position surrounding blood vessels and nerves in the peritendon (Figure 2). Cell-cell communication analysis indicated a direct interaction between NEGR1hi VCANhi fibroblasts and vascular endothelial cells via *SCARA5-VWR*, so they likely have a structural role supporting tendon vascularisation and innervation. Fibroblast-enabled neovascularisation is a hallmark of Achilles tendinopathy (30), so the role of NEGR1hi VCANhi fibroblasts in disease pathogenesis remains an important topic for future study.

NEGR1hi ITGA6hi fibroblasts were observed across the whole tendon-muscle unit and were spatially located surrounding peripheral nerve bundles. They expressed genes related to cell junctions and neurons/axons and interacted directly with Schwann cells via *NRXN1-NLGN1*. These fibroblasts had similar gene expression patterns to the hamstring tendon “nerve cells” (*TENM2, SLC22A3*) (14), quadriceps tendon ABCA10+ fibroblasts (*ACBA10, CNTN4*) (13) and also to endo-and perineurial fibroblasts found in skeletal muscle (*ITGA6, SLC22A3*) (31), so are likely nerve-associated or perineural fibroblasts. Since injuries to the tendon-muscle unit are a significant health issue (32, 33), the role of these cells should be of special interest in future studies in terms of prevention and modulation of pain in tendon pathologies.

COMPhi THBS4hi fibroblasts and NEGR1hi COL15A1hi fibroblasts were mainly detected in MTJ, or in muscle and MTJ, respectively. Both these fibroblast subsets were enriched for muscle-related processes and the *MYF6* regulon, a myogenic transcription factor, suggesting that these fibroblast types support the formation of muscle fibres. NEGR1hi COL15A1hi fibroblasts had a similar transcriptional profile to parenchymal fibroblasts in muscle (*COL15A1, COL4A1, ALDH1A2*) (18) as well as “universal” fibroblasts found across multiple tissues (*COL15A1, COL4A1, HSPG2*) (20, 21), indicating that they are likely a common fibroblast present in muscle and other non-tendon tissues. Since *THBS4* was abundant in fibroblasts from mouse Achilles MTJ (18) and hamstring MTJ (22), the COMPhi THBS4hi fibroblasts identified in our single-nuclei sequencing are likely an MTJ-specific subset.

A small number of *PRG4hi* fibroblasts were detected in the snRNA-seq data, predominantly in the tendon enthesis and midbody zones. The key marker *PRG4* (lubricin) facilitates biological lubrication, for example as part of the synovial fluid. While mainly known as a key marker for lining layer fibroblasts in synovium (34, 35), expression of *PRG4* has previously been shown in fibrocartilaginous region of tendon (36) and has been proposed to be important for surface lubrication of the tendon sheath (37).

Chondrocytes were defined by their expression of canonical markers (*ACAN, COL2A1*, *WWP2*), and enrichment of the SOX9 regulon, a known regulator of chondrocytes. These cells show similarity to the chondrocytes found in supraspinatus tendon (20) and pre-hypertrophic chondrocytes in cartilage (38). The presence of these cells specifically in tendon enthesis zone single-nuclei data is in accordance with the presence of collagen II and aggrecan in enthesis, but not midbody of tendon (39).

Over 20 (sub)types of non-fibroblast cells were identified. Apart from skeletal muscle cell types that were highly enriched in the MTJ and muscle regions, few cell types showed strong regional changes like those found with fibroblast subsets. This supports the hypothesis that tendon fibroblasts are not only critical for the construction and maintenance of tendon as key producers of ECM but also support the establishment of specialised tissue niches through cell-cell interaction of varying fibroblast subsets with other resident cell types. The existence of the proposed specialised fibroblast subsets in health raises the question of how these different subsets are affected by different pathologies. The Achilles is the most heavily mechanically loaded tendon in the human body and the most frequently injured, with consequent burden on society and healthcare systems. Previous studies have shown that genetic variants at *COL15A1* and *MMP3* loci, two genes highly expressed in different fibroblast subsets, are associated with Achilles tendinopathy (28–30), implicating the specific fibroblast subsets expressing these genes with the disease. Future elucidation of the cellular basis of pathogenesis of Achilles tendinopathies, including assessment of microanatomical differences in disease susceptibility, will lead to long-term improvements in patient care.

Healthy tendon tissue is challenging to acquire for research making this dataset a valuable resource. In this study we obtained healthy tendon tissue from patients undergoing an amputation due to sarcoma. The Achilles tendons all looked macroscopically normal, came from donors without a history of tendon disease, and donors were ambulant at the time of surgery.

Spatial transcriptomics was able to confirm the presence and location of most cell types, thereby adding a wealth of knowledge on specialised cell types and functions across microanatomy. However, the location of a few small (sub-) populations of cells could not be confirmed due to limitations of available tissue sections, as well as a limited number of gene probes (480) used. For example, chondrocytes have been hypothesized to be present in small numbers, only forming 3-4 rows of cells at the bone-tendon interface (10), so while they were detected in snRNA-seq data from the tendon enthesis zone, we were unable to confirm their location in the spatial data. Similarly, muscle fibres and corresponding associated fibroblasts were not captured in the MTJ spatial section.

In conclusion, this transcriptomic atlas of healthy human Achilles tendon describes the cellular composition of the tendon at unprecedented resolution, which has been vital to progress our understanding of the transitional nature of the bone-tendon-muscle unit. The changes in the presence and proportion of cell types, especially fibroblast subsets, across the different microanatomical zones of the Achilles tendon-muscle unit suggest functional niches that might be uniquely disrupted by injuries or autoimmune diseases. Therefore, this work forms a critical reference dataset for future studies on both tendon development and repair, and the pathologies affecting the different regions of the Achilles tendon-muscle unit.

## MATERIALS AND METHODS

### Ethics

Ethical approval was granted for the Oxford Musculoskeletal Biobank (19/SC/0134) by the local research ethics committee (Oxford Research Ethics Committee B) for all work on Achilles tendon, and informed written consent was obtained from all patients according to the Declaration of Helsinki.

### Tissue acquisition and processing

Achilles tendon tissue was collected from patients undergoing above knee lower leg amputation due to non-reconstructable sarcoma. Patients were excluded from the study if they had an active infection at time of surgery or were not weight-bearing. The age, sex, and affected side of the donors are reported alongside the de-identified study ID of each patient (Table 1). Tendon tissue was processed as previously described (40). Briefly, tissue was collected in cold DMEM-F12 media (Gibco) supplemented with 10% Foetal Bovine Serum (Labtech International) and 1% Penicillin/Streptomycin (Gibco). The collected sample was processed within two hours of tissue collection. Tendon was washed in PBS, fat and muscle were dissected off, and the tendon was cut into 1 cm tissue strips from the enthesis to MTJ; the 1-cm wide tissue strips were then cut into 1 cm pieces. Tissues were photographed to retain topographical reference. Adjoining muscle tissue (triceps surae) was collected from three patients (Table 1). The pieces of tendon and muscle tissue were snap-frozen in cryotubes using liquid nitrogen and stored at - 80°C until use. Some pieces of tendon and muscle were fixed by immersing the tissues in 10% formalin for 0.5 mm/hour at room temperature , washed, and transferred to 70% Industrial Methylated Spirits (IMS) for short term storage at 4°C until wax embedding.

### H&E imaging

Paraffin-embedded Achilles samples were sectioned at a thickness of 5μm, deparaffinised and rehydrated. They were incubated with Gill’s III Haematoxylin (Sigma-Aldrich) for 8 minutes, blued with 0.1% sodium bicarbonate solution for 30 seconds, and differentiated with 0.3% acid alcohol for 30 seconds. The sections were then stained with 0.5% Aqueous Eosin Y solution (Sigma-Aldrich) acidified with glacial acetic acid for 5 minutes and the excess stain was washed off. The slides were then dehydrated, mounted with DPX (Sigma-Aldrich), and imaged with the MoticEasyScan One digital slide scanner (RRID:SCR_024855).

### Single nucleus RNA sequencing

#### Nuclei isolation

Nuclei were isolated from the snap-frozen tissue using our previously published protocol (41). In short, Achilles tendon was cut into thin sections (from tendon tissue from proximal to enthesis, midbody, or proximal to the MTJ) using a scalpel, forceps, and a petri-dish that were all pre-cooled on dry ice and stored in a 50mL Falcon tube at -80°C until use. Adjacent triceps surae muscle was cut similarly. On the day of cell lysis, the tubes containing the cut tissue were thawed, and 4 mL of cold 1X CST buffer (containing NaCl, Tris-HCl pH 7.5, CaCl2, and MgCl2, with CHAPS hydrate (Sigma), BSA (Sigma), RNase inhibitors (RNaseIn Plus; Promega and SUPERase In; Invitrogen), and protease inhibitor (cOmplete tablet; Roche); full recipe can be found in the published protocol) was added. After 10 minutes of incubation on a rotor at 4°C, the tissue/buffer mixture was poured through a 40 µm strainer and the tube used for tissue lysis was washed twice with 2mL PBS with 1% BSA. The nuclei solution was then transferred to a 15mL Falcon tube, and the previous 50mL tube was washed once with 4 mL PBS with 1% BSA and transferred to the same 15mL Falcon tube. The nuclei solution was centrifuged at 500g at 4°C for 5 minutes. After pouring off the supernatant, the tubes were briefly spun down, the nuclei were resuspended in the supernatant, and the remaining volume was determined. The concentration of nuclei was determined by counting DAPI-stained nuclei in a Neubauer Improved haemocytometer (NanoEnTek).

#### Library preparation and sequencing

Nuclei suspensions were diluted (PBS with 1% BSA) to 860-1,000 nuclei/µl and loaded onto the 10X Genomics Chromium Controller Genetic Analyzer (RRID:SCR_019326), then libraries were prepared using the Chromium Next GEM Single Cell 3’ Reagent Kits v3.1 (10x Genomics) following the manufacturer’s instructions and indexed with the Single Index Kit T Set A (10x Genomics). Quality control of cDNA and final libraries was analysed using High Sensitivity ScreenTape assays on Agilent 4150 TapeStation System (RRID:SCR_019393). Final libraries were pooled and sequenced on an Illumina NovaSeq 6000 Sequencing System (RRID:SCR_016387) by Azenta Life Sciences at a minimum depth of ∼20,000 read pairs per expected nuclei.

### Single nucleus RNA sequencing data analysis

#### Quality control and filtering

Raw NGS data was quality controlled and mapped using the *scflow quantnuclei* pipeline (https://github.com/cribbslab/scflow; python v3.8.15). Specifically, quality control of Fastq files was performed using fastqc (v0.12.1). Fastq files were mapped to the human Ensembl GRCh38 transcriptome (release 106) using kb_python (RRID:SCR_018213) command kb count (v0.27.3) with kmer size = 31. Spliced and unspliced matrices were merged to create the count matrix for downstream analysis. The snRNA-seq analysis was performed according to best practises (42), using R Project for Statistical Computing (RRID:SCR_001905) (v4.4.1) and RStudio (RRID:SCR_000432) Server (v2024.09.0 build 375) with SingleCellExperiment (RRID:SCR_026794) (43) (v1.24.0) and Seurat (RRID:SCR_016341) (44) (v4.3.0.1) tools, and mamba (https://github.com/mamba-org/mamba) (v1.4.4) for version control. Poor quality cells were removed by filtering on a per-sample basis for low number of cells and features and high mitochondrial ratio. Ambient RNA was detected using the decontX function from celda (v1.14.0)(45) and droplets with decontX score > 0.3 were removed. Genes with >2% ambient contamination were removed (8 genes). Doublets were calculated and removed with scDblFinder(46) (v1.12.0). Ambient RNA was then detected again using SoupX (RRID:SCR_019193) (47) (v1.6.2) using parameters customised to each sample. Highly contaminated droplets (>30% soupX contamination) were removed and the soupX adjusted count matrix was used for downstream analysis.

#### Data integration, clustering and broad annotation

Using Seurat all datasets were merged, log-normalised, 5,000 variable features were selected, the data was scaled, and 50 PCs were calculated. A custom script was used to convert the Seurat object to Anndata (RRID:SCR_018209) format. Integration across patient and sequencing batch was performed using scvi-tools (RRID:SCR_026673) (48) (v1.0.2) using Jupyter Notebook (RRID:SCR_018315) (v7.2.1). Ray autotune was used to select optimal hyperparameters which were then used in the scVI model. The model was trained using vae.train for 60 epochs then latent representation embeddings were extracted and returned to the Seurat object for downstream visualisation and analysis. Calculation of neighbours, Louvain clustering, and dimensionality reduction were then performed in Seurat using the scVI embeddings and 40 PCs. Integration success was measured using elbow plots, and calculation of LISI scores (LISI v1.0) (49). Clustree (RRID:SCR_016293) (v0.5.0) was used to assess optimal clustering resolution (50). Each cluster comprised cells from at least three patients and no one cluster had >80% cells from one patient, with the exception of B cells which were mostly from patient MSK1687 (96.8%). Specific marker genes were calculated using Seurat FindAllMarkers() and visualised using Seurat DoHeatmap() and scDotPlot (v1.1.0). Broad cell-type annotations were assigned according to expression of canonical marker genes described in the literature visualised with Seurat DotPlot(). SCpubr (v2.0.2) was used for improved visualisation (51).

#### Fine annotation

Each broad cell type comprising fibroblasts, muscle cells (skeletal muscle cells and satellite cells), immune cells (macrophages, T cells, B cells, plasma cells, granulocytes) and non-fibroblast stromal cells (adipocytes, lymphatic endothelial cells, vascular endothelial cells, Schwann cells, mural cells) was then subset and re-clustered using the methods described above. Crucially this allowed feature selection (2000 variable genes) and dimensionality reduction to be specific to each cell subset, revealing additional distinct cell types that were defined according to literature markers and/or highly expressed genes. Integration of the muscle subset with scVI failed quality control as several clusters contained >80% of cells from one patient so in this case reintegration was performed with Harmony (RRID:SCR_022206) (49) (v1.2.0). In some cases, smaller clusters were merged if they did not express biologically meaningful distinct marker genes.

#### Differential abundance of cell types across microanatomy

MiloR (RRID:SCR_025630)(52) (v1.10.0) was used to assess differential abundance of broad cell types or fibroblasts across microanatomy using broad or fine annotations.

#### Differential gene expression and pathway enrichment

To determine differential gene expression (DGE) within each broad cell type across microanatomy, gene expression was pseudobulked per cell type using Seurat AggregateExpression(), with genes with less than 10 reads filtered out. DESeq2 (RRID:SCR_015687) (53) (v1.40.2) was used to identify differentially expressed genes across microanatomy using the design formula “∼ patient + microanatomical_site”, and the likelihood ratio test (LRT) used to compare multiple conditions. Pre-and post-QC heatmaps and PCA plots were generated, and comparisons were only drawn where cells from at least three donors were present at each microanatomy. Differentially expressed genes were clustered and visualised using DEGreport (RRID:SCR_018941) (v1.36.0). ClusterProfiler (RRID:SCR_016884) (54) (v4.10.0) was used to obtain enriched GO:BP pathways on genes from groups of clusters into the following defined classes: enthesis-high, muscle-high, tendon-high. A custom background of all genes expressed in that cell type was given and results were plotted using ggplot2 (RRID:SCR_014601). The same methods were also used to assess pathways enriched in each fibroblast subtype. Gene expression in each fibroblast subtype was aggregated by pseudobulk and the top 200 genes by number of counts was as the input to clusterProfiler.

#### Ligand-receptor interactions

Liana (55) (v0.1.14) was used to identify ligand-receptor interactions between finely annotated cell types at each microanatomy, using CellPhoneDB (RRID:SCR_017054) and selecting interactions with a p-value of at least 0.05. CrossTalkeR (56) (v1.4.0) was used to visualise overall ligand-receptor interactions using plot_cci(), and liana_dotplot was used to visualise specific interactions between selected cell types.

### Spatial transcriptomics

#### Sample preparation and 10X Xenium

Representative tissue samples of the three tendon microanatomies and adjacent muscle were selected based on initial haematoxylin and eosin (H&E) staining. Regions of interest (ROIs) were identified and embedded into a Tissue Array measuring 10.45 mm × 22.45 mm, corresponding to the dimensions of the Sample Area on 10X Genomics Xenium slides. Sectioning was performed as per Xenium “In Situ-Tissue Preparation Guide protocol” (10X Genomics, Demonstrated Protocol, CG000578 Rev E). 5μm sections were cut and briefly placed in Milli-Q water at 37°C then transferred to the Xenium slide (PN-3000941). Slides were kept upright at 40°C until excess water had dripped off and evaporated, then baked at 45°C for 3 hours. The slides were stored in a desiccator at room temperature until they were ready for Xenium analysis. Slides underwent deparaffinisation and decrosslinking according to the 10X Xenium “In Situ for FFPE-Deparaffinization and Decrosslinking Protocol” (10X Genomics, CG000580 Rev E) and using the Xenium Slides and Sample Prep Reagents Kit (10X Genomics).

Hybridisation was performed according to the Xenium In Situ Gene Expression User Guide (10X Genomics, CG000749 Rev B) C with a Probe Hybridization Mix containing a bespoke Add-On Custom Probe Panel (gene list in Data S3) of 101 genes designed to differentiate cell types found in the scRNA-seq data, and the Xenium Human Multi Tissue and Cancer Probe Panel of 300 genes. Standard protocols were followed for cell segmentation staining and analysis was performed using a Xenium Analyzer (RRID:SCR_023910) according to the Xenium Analyzer User Guide (10X Genomics, CG000584 Rev J). Following analysis, slides were stored at 4°C under PBS-Tween.

#### Post-Xenium H&E staining

H&E staining was performed according to Xenium’s Post-Xenium Analyzer H&E Staining (10X Genomics, CG000613 Rev A) protocol. Coverslips were mounted on the slides with DPX and the slides were imaged with MoticEasyScan One digital slide scanner (RRID:SCR_024855).

### Spatial transcriptomics data analysis

Xenium data was processed using the 10X Genomics Xenium Onboard Analysis (RRID: SCR_026158). Output files were analysed using Seurat (RRID:SCR_016341) v5.3.0 in R (RRID:SCR_001905) v4.5.1 with RStudio (RRID:SCR_000432) Server Pro build 467.pro1. Filtering was performed according to cell size, number of transcripts and number of probes. Data from all four tissue pieces was integrated using Harmony (RRID:SCR_022206) with Seurat IntegrateLayers(). Normalisation was performed using SCTransform (RRID:SCR_022146), and the Seurat workflow was followed for dimensionality reduction (RunPCA(), RunUMAP()).Annotation was performed by transferring broad and fine annotation labels from the snRNA-seq data using FindTransferAnchors() and TransferData(). Broad cell types (immune, fibroblast, muscle and stromal) were subsetted, and the above steps were repeated to define the finely annotated cell types. Annotations were uploaded to 10x Genomics Xenium Explorer (RRID:SCR_025847) along with post-Xenium H&E staining to generate figures. SCpubr_2.0.2 was used for UMAP plots.

## Acknowledgements

Funding: This work was funded by the following grants:

- Chan Zuckerberg Initiative Tendon Seed Network grant Chan Zuckerberg Initiative (CZI). Grant Numbers: 2019-002426, 2021-240342
- Versus Arthritis. Grant Number: 22873
- Rhodes Scholarships (The Rhodes Scholarship)
- Oxford-Medical Research Council Doctoral Training Partnership
- NIHR | NIHR Oxford Biomedical Research Centre (OxBRC)
- Medical Research Council. Grant Number: MR/V010182/1
- Paget’s Association. Grant Number: PA21010

We thank our research assistant Louise Appleton, our research nurses Debra Beazley and Lois Vesty-Edwards, and the surgical teams at the Nuffield Orthopaedic Centre for their invaluable help collecting human tissue for this study. We would like acknowledge support from the Oxford Musculoskeletal Biobank and thank the patients for donating their tissue to research. We thank Joanna Hester and Audrey Au Young for assistance running the Xenium experiments. Finally, we thank all other members of the CZI Tendon Seed Network for insightful discussions.

## Author Contributions

Conceptualization: APC, MJB, SJBS

Data Curation: CJC, JYM, APC, KRA

Formal analysis: CJC, AK, KRA

Funding acquisition: APC, MJB, SJBS

Investigation: JYM, AK, CP, SS, LRM, CTI, ACA, TBS, MN

Resources: CG, DW, TC, SG, AS, RBR, HBW, APC, DS, MJB, SJBS

Supervision: PH, DS, APC, MJB, SJBS

Visualization: CJC

Writing – Original draft: CJC

Writing – Review & Editing: CJC, JYM, PH, DS, APC, MJB, SJBS

## Competing Interests

APC is a cofounder and employee of Caeruleus Genomics Ltd and is an inventor on several patents related to sequencing technologies filed by Oxford University Innovations. All other authors declare they have no competing interests.

Data and materials availability: All the code used for the data analysis of this paper will be made available on GitHub. All the transcriptomics data in this manuscript will be made available upon publication.

## REFERENCES

1. M. N. Doral, M. Alam, M. Bozkurt, E. Turhan, O. A. Atay, G. Dönmez, N. Maffulli, Functional anatomy of the Achilles tendon. Knee Surgery, Sports Traumatology, Arthroscopy 18, 638–643 (2010).

2. S. A. Hefferan, C. L. Blaker, D. M. Ashton, C. B. Little, E. C. Clarke, Structural Variations of Tendons: A Systematic Search and Narrative Review of Histological Differences Between Tendons, Tendon Regions, Sex, and Age. Journal of Orthopaedic Research.

3. P. W. Ackermann, P. Phisitkul, C. J. Pearce, Achilles tendinopathy – pathophysiology: state of the art. Journal of ISAKOS 3, 304–314 (2018).

4. A. Traweger, A. Scott, M. Kjaer, E. Wezenbeek, R. Scattone Silva, J. G. Kennedy, J. J. Butler, M. Gomez-Florit, M. E. Gomes, J. G. Snedeker, S. G. Dakin, B. Wildemann, Achilles tendinopathy. Nature Reviews Disease Primers 11, 20 (2025).

5. A. Alvarez, T. K. Tiu, Enthesopathies in StatPearls. (StatPearls Publishing, 2025, Treasure Island (FL), 2025).

6. M. Baldwin, C. D. Buckley, F. Guilak, P. Hulley, A. P. Cribbs, S. Snelling, A roadmap for delivering a human musculoskeletal cell atlas. Nat Rev Rheumatol 19, 738–752 (2023).

7. K. Winnicki, A. Ochała-Kłos, B. Rutowicz, P. A. Pękala, K. A. Tomaszewski, Functional anatomy, histology and biomechanics of the human Achilles tendon — A comprehensive review. Annals of Anatomy - Anatomischer Anzeiger 229, 151461 (2020).

8. I. Tresoldi, F. Oliva, M. Benvenuto, M. Fantini, L. Masuelli, R. Bei, A. Modesti, Tendon’s ultrastructure. Muscles Ligaments Tendons J 3, 2–6 (2013).

9. J. R. Jakobsen, M. R. Krogsgaard, The Myotendinous Junction—A Vulnerable Companion in Sports. A Narrative Review. Frontiers in Physiology 12, (2021).

10. J. Apostolakos, T. J. Durant, C. R. Dwyer, R. P. Russell, J. H. Weinreb, F. Alaee, K. Beitzel, M. B. McCarthy, M. P. Cote, A. D. Mazzocca, The enthesis: a review of the tendon-to-bone insertion. Muscles Ligaments Tendons J 4, 333–342 (2014).

11. R. Elmentaite, N. Kumasaka, K. Roberts, A. Fleming, E. Dann, H. W. King, V. Kleshchevnikov, M. Dabrowska, S. Pritchard, L. Bolt, S. F. Vieira, L. Mamanova, N. Huang, F. Perrone, I. Goh Kai’En, S. N. Lisgo, M. Katan, S. Leonard, T. R. W. Oliver, C. E. Hook, K. Nayak, L. S. Campos, C. Domínguez Conde, E. Stephenson, J. Engelbert, R. A. Botting, K. Polanski, S. van Dongen, M. Patel, M. D. Morgan, J. C. Marioni, O. A. Bayraktar, K. B. Meyer, X. He, R. A. Barker, H. H. Uhlig, K. T. Mahbubani, K. Saeb-Parsy, M. Zilbauer, M. R. Clatworthy, M. Haniffa, K. R. James, S. A. Teichmann, Cells of the human intestinal tract mapped across space and time. Nature 597, 250–255 (2021).

12. B. B. Lake, R. Menon, S. Winfree, Q. Hu, R. Melo Ferreira, K. Kalhor, D. Barwinska, E. A. Otto, M. Ferkowicz, D. Diep, N. Plongthongkum, A. Knoten, S. Urata, L. H. Mariani, A. S. Naik, S. Eddy, B. Zhang, Y. Wu, D. Salamon, J. C. Williams, X. Wang, K. S. Balderrama, P. J. Hoover, E. Murray, J. L. Marshall, T. Noel, A. Vijayan, A. Hartman, F. Chen, S. S. Waikar, S. E. Rosas, F. P. Wilson, P. M. Palevsky, K. Kiryluk, J. R. Sedor, R. D. Toto, C. R. Parikh, E. H. Kim, R. Satija, A. Greka, E. Z. Macosko, P. V. Kharchenko, J. P. Gaut, J. B. Hodgin, R. Knight, S. H. Lecker, I. Stillman, A. A. Amodu, T. Ilori, S. Maikhor, I. Schmidt, G. M. McMahon, A. Weins, N. Hacohen, L. Bush, A. Gonzalez-Vicente, J. Taliercio, J. O’toole, E. Poggio, L. Cooperman, S. Jolly, L. Herlitz, J. Nguyen, E. Palmer, D. Sendrey, K. Spates-Harden, P. Appelbaum, J. M. Barasch, A. S. Bomback, V. D. D’Agati, K. Mehl, P. A. Canetta, N. Shang, O. Balderes, S. Kudose, L. Barisoni, T. Alexandrov, Y. Cheng, K. W. Dunn, K. J. Kelly, T. A. Sutton, Y. Wen, C. P. Corona-Villalobos, S. Menez, A. Rosenberg, M. Atta, C. Johansen, J. Sun, N. Roy, M. Williams, E. U. Azeloglu, C. He, R. Iyengar, J. Hansen, Y. Xiong, B. Rovin, S. Parikh, S. M. Madhavan, C. R. Anderton, L. Pasa-Tolic, D. Velickovic, O. Troyanskaya, R. Sealfon, K. R. Tuttle, Z. G. Laszik, G. Nolan, M. Sarwal, K. Anjani, T. Sigdel, H. Ascani, U. G. J. Balis, C. Lienczewski, B. Steck, Y. He, J. Schaub, V. M. Blanc, R. Murugan, P. Randhawa, M. Rosengart, M. Tublin, T. Vita, J. A. Kellum, D. E. Hall, M. M. Elder, J. Winters, M. Gilliam, C. E. Alpers, K. N. Blank, J. Carson, I. H. De Boer, A. L. Dighe, J. Himmelfarb, S. D. Mooney, S. Shankland, K. Williams, C. Park, F. Dowd, R. L. McClelland, S. Daniel, A. N. Hoofnagle, A. Wilcox, S. Bansal, K. Sharma, M. Venkatachalam, G. Zhang, A. Pamreddy, V. R. Kakade, D. Moledina, M. M. Shaw, U. Ugwuowo, T. Arora, J. Ardayfio, J. Bebiak, K. Brown, C. E. Campbell, J. Saul, A. Shpigel, C. Stutzke, R. Koewler, T. Campbell, L. Hayashi, N. Jefferson, R. Pinkeney, G. V. Roberts, M. T. Eadon, P. C. Dagher, T. M. El-Achkar, K. Zhang, M. Kretzler, S. Jain, K. Consortium, An atlas of healthy and injured cell states and niches in the human kidney. Nature 619, 585–594 (2023).

13. J. Y. Mimpen, M. J. Baldwin, C. Paul, L. Ramos-Mucci, A. Kurjan, C. J. Cohen, S. Sharma, M. S. N. Chevalier Florquin, P. A. Hulley, J. McMaster, A. Titchener, A. Martin, M. L. Costa, S. E. Gwilym, A. P. Cribbs, S. J. B. Snelling, Exploring cellular changes in ruptured human quadriceps tendons at single-cell resolution. *bioRxiv*, 2024.2009.2006.611599 (2024).

14. J. Y. Mimpen, L. Ramos-Mucci, C. Paul, A. Kurjan, P. A. Hulley, C. T. Ikwuanusi, C. J. Cohen, S. E. Gwilym, M. J. Baldwin, A. P. Cribbs, S. J. B. Snelling, Single nucleus and spatial transcriptomic profiling of healthy human hamstring tendon. Faseb j 38, e23629 (2024).

15. A. R. Kendal, T. Layton, H. Al-Mossawi, L. Appleton, S. Dakin, R. Brown, C. Loizou, M. Rogers, R. Sharp, A. Carr, Multi-omic single cell analysis resolves novel stromal cell populations in healthy and diseased human tendon. Scientific Reports 10, 13939 (2020).

16. M. Akbar, L. MacDonald, L. A. N. Crowe, K. Carlberg, M. Kurowska-Stolarska, P. L. Ståhl, S. J. B. Snelling, I. B. McInnes, N. L. Millar, Single cell and spatial transcriptomics in human tendon disease indicate dysregulated immune homeostasis. Annals of the Rheumatic Diseases 80, 1494–1497 (2021).

17. F. Fang, Y. Xiao, E. Zelzer, K. W. Leong, S. Thomopoulos, A mineralizing pool of Gli1-expressing progenitors builds the tendon enthesis and demonstrates therapeutic potential. Cell Stem Cell 29, 1669–1684.e1666 (2022).

18. R. Yan, H. Zhang, Y. Ma, R. Lin, B. Zhou, T. Zhang, C. Fan, Y. Zhang, Z. Wang, T. Fang, Z. Yin, Y. Cai, H. Ouyang, X. Chen, Discovery of Muscle-Tendon Progenitor Subpopulation in Human Myotendinous Junction at Single-Cell Resolution. Research (Wash D C) 2022, 9760390 (2022).

19. T. Zhang, L. Wan, H. Xiao, L. Wang, J. Hu, H. Lu, Single-cell RNA sequencing reveals cellular and molecular heterogeneity in fibrocartilaginous enthesis formation. eLife 12, e85873 (2023).

20. W. Fu, R. Yang, J. Li, Single-cell and spatial transcriptomics reveal changes in cell heterogeneity during progression of human tendinopathy. BMC Biology 21, 132 (2023).

21. A. Kurjan, J. Y. Mimpen, L. Ramos-Mucci, A. C. Aksu, C. D. Buckley, A. P. Cribbs, M. J. Baldwin, S. J. B. Snelling, Cellular and molecular landscapes of human tendons across the lifespan revealed by spatial and single-cell transcriptomics. bioRxiv, 2025.2006.2019.660575 (2025).

22. A. Møbjerg, D. Steffen, P. Schjerling, J. R. Jakobsen, A. Jokipii-Utzon, M. Y. Batiuk, K. Khodosevich, M. R. Krogsgaard, V. Izzi, A. L. Mackey, M. Kjaer, C.-Y. C. Yeung, Spatially distinct ECM-producing fibroblasts and myonuclei orchestrate early adaptation to mechanical loading in the human muscle-tendon unit. *bioRxiv*, 2025.2008.2028.672815 (2025).

23. A. J. De Micheli, J. B. Swanson, N. P. Disser, L. M. Martinez, N. R. Walker, D. J. Oliver, B. D. Cosgrove, C. L. Mendias, Single-cell transcriptomic analysis identifies extensive heterogeneity in the cellular composition of mouse Achilles tendons. American Journal of Physiology-Cell Physiology 319, C885–C894 (2020).

24. C. Bravo González-Blas, S. De Winter, G. Hulselmans, N. Hecker, I. Matetovici, V. Christiaens, S. Poovathingal, J. Wouters, S. Aibar, S. Aerts, SCENIC+: single-cell multiomic inference of enhancers and gene regulatory networks. Nature Methods 20, 1355–1367 (2023).

25. Y. Zhang, L. Zhang, X.-h. Lin, Z.-m. Li, Q.-y. Zhang, Knockdown of IRF3 inhibits extracellular matrix expression in keloid fibroblasts. Biomedicine & Pharmacotherapy 88, 1064–1068 (2017).

26. F. Lazure, D. M. Blackburn, A. H. Corchado, K. Sahinyan, N. Karam, A. Sharanek, D. Nguyen, C. Lepper, H. S. Najafabadi, T. J. Perkins, A. Jahani-Asl, V. D. Soleimani, Myf6/MRF4 is a myogenic niche regulator required for the maintenance of the muscle stem cell pool. EMBO Rep 21, e49499 (2020).

27. M. Kurowska-Stolarska, S. Alivernini, Synovial tissue macrophages in joint homeostasis, rheumatoid arthritis and disease remission. Nature Reviews Rheumatology 18, 384–397 (2022).

28. A. Karlsen, C.-Y. C. Yeung, P. Schjerling, L. Denz, C. Hoegsbjerg, J. R. Jakobsen, M. R. Krogsgaard, M. Koch, S. Schiaffino, M. Kjaer, A. L. Mackey, Distinct myofibre domains of the human myotendinous junction revealed by single-nucleus RNA sequencing. Journal of Cell Science 136, (2023).

29. Y. Asano, M. Trojanowska, Fli1 Represses Transcription of the Human α2(I) Collagen Gene by Recruitment of the HDAC1/p300 Complex. PLOS ONE 8, e74930 (2013).

30. S. G. Dakin, J. Newton, F. O. Martinez, R. Hedley, S. Gwilym, N. Jones, H. A. B. Reid, S. Wood, G. Wells, L. Appleton, K. Wheway, B. Watkins, A. J. Carr, Chronic inflammation is a feature of Achilles tendinopathy and rupture. British Journal of Sports Medicine 52, 359–367 (2018).

31. V. R. Kedlian, Y. Wang, T. Liu, X. Chen, L. Bolt, C. Tudor, Z. Shen, E. S. Fasouli, E. Prigmore, V. Kleshchevnikov, J. P. Pett, T. Li, J. E. G. Lawrence, S. Perera, M. Prete, N. Huang, Q. Guo, X. Zeng, L. Yang, K. Polański, N.-J. Chipampe, M. Dabrowska, X. Li, O. A. Bayraktar, M. Patel, N. Kumasaka, K. T. Mahbubani, A. P. Xiang, K. B. Meyer, K. Saeb-Parsy, S. A. Teichmann, H. Zhang, Human skeletal muscle aging atlas. Nature Aging 4, 727–744 (2024).

32. A. S. Tadros, B. K. Huang, M. N. Pathria, Muscle-Tendon-Enthesis Unit. Semin Musculoskelet Radiol 22, 263–274 (2018).

33. A. G. Shamrock, M. A. Dreyer, M. A. Varacallo, Achilles Tendon Rupture in StatPearls. (StatPearls Publishing, 2025, Treasure Island (FL), 2025).

34. Y. Gao, J. Li, W. Cheng, T. Diao, H. Liu, Y. Bo, C. Liu, W. Zhou, M. Chen, Y. Zhang, Z. Liu, W. Han, R. Chen, J. Peng, L. Zhu, W. Hou, Z. Zhang, Cross-tissue human fibroblast atlas reveals myofibroblast subtypes with distinct roles in immune modulation. Cancer Cell 42, 1764–1783.e1710 (2024).

35. M. H. Smith, V. R. Gao, P. K. Periyakoil, A. Kochen, E. F. DiCarlo, S. M. Goodman, T. M. Norman, L. T. Donlin, C. S. Leslie, A. Y. Rudensky, Drivers of heterogeneity in synovial fibroblasts in rheumatoid arthritis. Nature Immunology 24, 1200–1210 (2023).

36. S. G. Rees, J. R. Davies, D. Tudor, C. R. Flannery, C. E. Hughes, C. M. Dent, B. Caterson, Immunolocalisation and expression of proteoglycan 4 (cartilage superficial zone proteoglycan) in tendon. Matrix Biol 21, 593–602 (2002).

37. D. K. Rhee, J. Marcelino, M. Baker, Y. Gong, P. Smits, V. Lefebvre, G. D. Jay, M. Stewart, H. Wang, M. L. Warman, J. D. Carpten, The secreted glycoprotein lubricin protects cartilage surfaces and inhibits synovial cell overgrowth. J Clin Invest 115, 622–631 (2005).

38. H. Li, X. Jiang, Y. Xiao, Y. Zhang, W. Zhang, M. Doherty, J. Nestor, C. Li, J. Ye, T. Sha, H. Lyu, J. Wei, C. Zeng, G. Lei, Combining single-cell RNA sequencing and population-based studies reveals hand osteoarthritis-associated chondrocyte subpopulations and pathways. Bone Research 11, 58 (2023).

39. A. D. Waggett, J. R. Ralphs, A. P. Kwan, D. Woodnutt, M. Benjamin, Characterization of collagens and proteoglycans at the insertion of the human Achilles tendon. Matrix Biol 16, 457–470 (1998).

40. J. Y. Mimpen, C. Paul, L. Ramos-Mucci, T. S. Network, A. P. Cribbs, M. J. Baldwin, S. J. B. Snelling, SOP for snap-freezing tissues. protocols.io, (2022).

41. J. Y. Mimpen, C. Paul, T. S. Network, A. P. Cribbs, S. J. B. Snelling, Nuclei isolation from snap-frozen tendon tissue for single nucleus RNA Sequencing. protocols.io, (2021).

42. L. Heumos, A. C. Schaar, C. Lance, A. Litinetskaya, F. Drost, L. Zappia, M. D. Lücken, D. C. Strobl, J. Henao, F. Curion, H. Aliee, M. Ansari, P. Badia-i-Mompel, M. Büttner, E. Dann, D. Dimitrov, L. Dony, A. Frishberg, D. He, S. Hediyeh-zadeh, L. Hetzel, I. L. Ibarra, M. G. Jones, M. Lotfollahi, L. D. Martens, C. L. Müller, M. Nitzan, J. Ostner, G. Palla, R. Patro, Z. Piran, C. Ramírez-Suástegui, J. Saez-Rodriguez, H. Sarkar, B. Schubert, L. Sikkema, A. Srivastava, J. Tanevski, I. Virshup, P. Weiler, H. B. Schiller, F. J. Theis, C. Single-cell Best Practices, Best practices for single-cell analysis across modalities. Nature Reviews Genetics 24, 550–572 (2023).

43. R. A. Amezquita, A. T. L. Lun, E. Becht, V. J. Carey, L. N. Carpp, L. Geistlinger, F. Marini, K. Rue-Albrecht, D. Risso, C. Soneson, L. Waldron, H. Pagès, M. L. Smith, W. Huber, M. Morgan, R. Gottardo, S. C. Hicks, Orchestrating single-cell analysis with Bioconductor. Nature Methods 17, 137–145 (2020).

44. Y. Hao, S. Hao, E. Andersen-Nissen, W. M. Mauck, S. Zheng, A. Butler, M. J. Lee, A. J. Wilk, C. Darby, M. Zager, P. Hoffman, M. Stoeckius, E. Papalexi, E. P. Mimitou, J. Jain, A. Srivastava, T. Stuart, L. M. Fleming, B. Yeung, A. J. Rogers, J. M. McElrath, C. A. Blish, R. Gottardo, P. Smibert, R. Satija, Integrated analysis of multimodal single-cell data. Cell 184, 3573–3587.e3529 (2021).

45. S. Yang, S. E. Corbett, Y. Koga, Z. Wang, W. E. Johnson, M. Yajima, J. D. Campbell, Decontamination of ambient RNA in single-cell RNA-seq with DecontX. Genome Biology 21, 57 (2020).

46. P. L. Germain, A. Lun, C. Garcia Meixide, W. Macnair, M. D. Robinson, Doublet identification in single-cell sequencing data using scDblFinder. F1000Res 10, 979 (2021).

47. M. D. Young, S. Behjati, SoupX removes ambient RNA contamination from droplet-based single-cell RNA sequencing data. Gigascience 9, (2020).

48. A. Gayoso, R. Lopez, G. Xing, P. Boyeau, V. Valiollah Pour Amiri, J. Hong, K. Wu, M. Jayasuriya, E. Mehlman, M. Langevin, Y. Liu, J. Samaran, G. Misrachi, A. Nazaret, O. Clivio, C. Xu, T. Ashuach, M. Gabitto, M. Lotfollahi, V. Svensson, E. da Veiga Beltrame, V. Kleshchevnikov, C. Talavera-López, L. Pachter, F. J. Theis, A. Streets, M. I. Jordan, J. Regier, N. Yosef, A Python library for probabilistic analysis of single-cell omics data. Nature Biotechnology 40, 163–166 (2022).

49. I. Korsunsky, N. Millard, J. Fan, K. Slowikowski, F. Zhang, K. Wei, Y. Baglaenko, M. Brenner, P.-r. Loh, S. Raychaudhuri, Fast, sensitive and accurate integration of single-cell data with Harmony. Nature Methods 16, 1289–1296 (2019).

50. L. Zappia, A. Oshlack, Clustering trees: a visualization for evaluating clusterings at multiple resolutions. GigaScience 7, (2018).

51. E. Blanco-Carmona, Generating publication ready visualizations for Single Cell transcriptomics using SCpubr. bioRxiv, 2022.2002.2028.482303 (2022).

52. E. Dann, N. C. Henderson, S. A. Teichmann, M. D. Morgan, J. C. Marioni, Differential abundance testing on single-cell data using k-nearest neighbor graphs. Nature Biotechnology 40, 245–253 (2022).

53. M. I. Love, W. Huber, S. Anders, Moderated estimation of fold change and dispersion for RNA-seq data with DESeq2. Genome Biology 15, 550 (2014).

54. G. Yu, L.-G. Wang, Y. Han, Q.-Y. He, clusterProfiler: an R Package for Comparing Biological Themes Among Gene Clusters. OMICS: A Journal of Integrative Biology 16, 284–287 (2012).

55. D. Dimitrov, D. Türei, M. Garrido-Rodriguez, P. L. Burmedi, J. S. Nagai, C. Boys, R. O. Ramirez Flores, H. Kim, B. Szalai, I. G. Costa, A. Valdeolivas, A. Dugourd, J. Saez-Rodriguez, Comparison of methods and resources for cell-cell communication inference from single-cell RNA-Seq data. Nature Communications 13, 3224 (2022).

56. J. S. Nagai, N. B. Leimkühler, M. T. Schaub, R. K. Schneider, I. G. Costa, CrossTalkeR: analysis and visualization of ligand–receptorne tworks. Bioinformatics 37, 4263–4265 (2021).

