## Supplementary Figures for "Fibroblast specialisation across microanatomy in a single-cell atlas of healthy human Achilles tendon"

Carla J. Cohen et al

**Supplementary Data**
Figures S1-S9
Data S1-S3

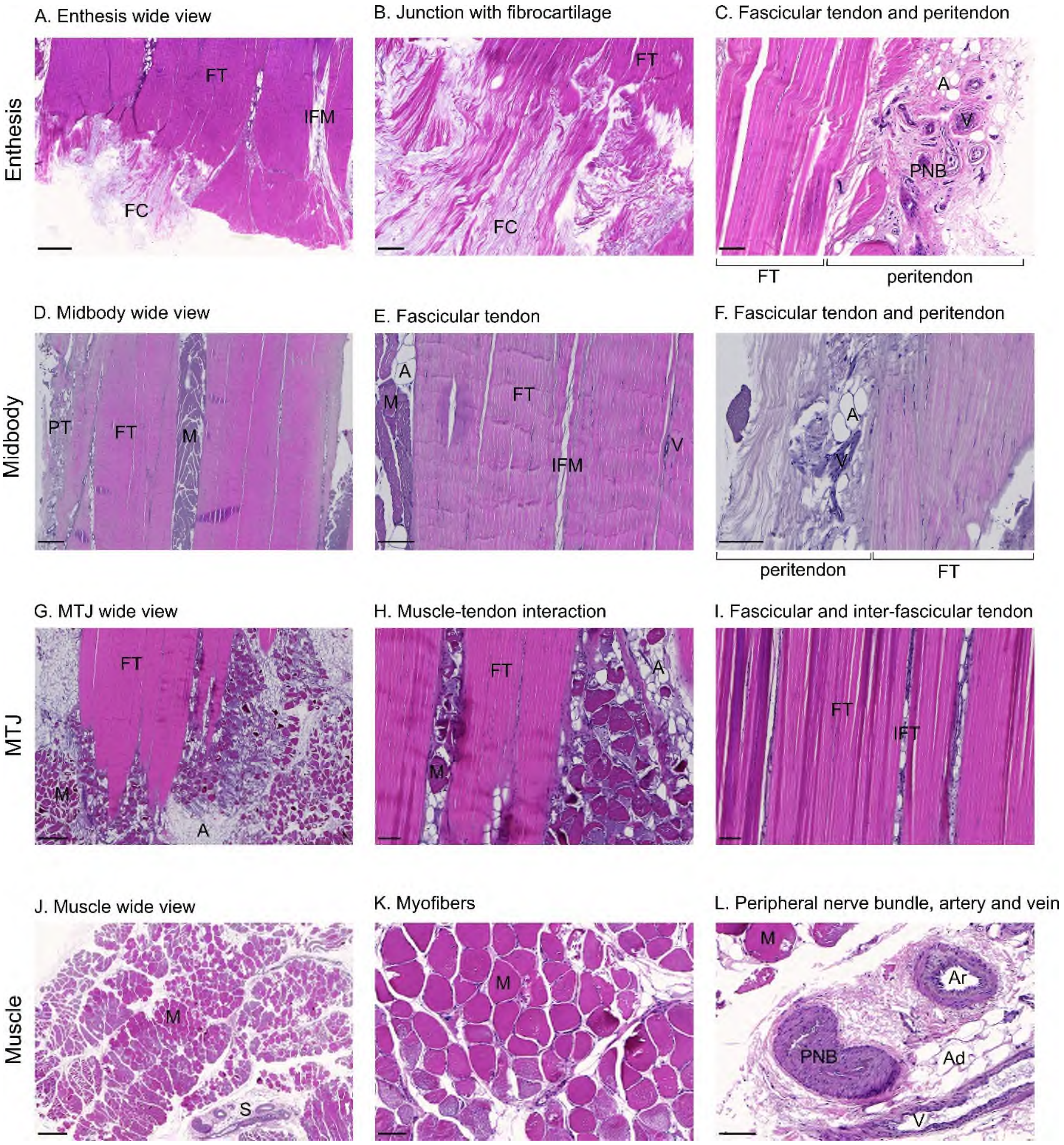
A. Enthesis wide view
B. Junction with fibrocartilage
C. Fascicular tendon and peritendon
Enthesis
FT
IFM
FC
A
V
PNB
peritendon
D. Midbody wide view
E. Fascicular tendon
F. Fascicular tendon and peritendon
Midbody
PT
M
IFM
G. MTJ wide view
H. Muscle-tendon interaction
I. Fascicular and inter-fascicular tendon
MTJ
IFT
J. Muscle wide view
K. Myofibers
L. Peripheral nerve bundle, artery and vein
Muscle
S
Ar
Ad

**Fig.S1. H&E images showing tendon and muscle architecture and tissue features**. Left panels are wide view with scale bar 500 µm; Middle and right panels are higher magnification with scale bar 100 µm. Tendon images are longitudinal cuts through the tissue; the muscle images are transverse. A-C are from the enthesis region. (A) Region surrounding the enthesis showing the collagen-rich striated fibrils of fascicular tendon (FT) joining with acellular fibrocartilage (FC), and interfascicular matrix (IFM). (B) Fascicular tendon junction with fibrocartilage (FC). (C) Fascicular tendon (FT) adjacent to the peritendon, which contains adipocytes (A), vessels (V) and peripheral nerve bundles (PNB). D-F are from the tendon midbody. (D) Wide view of tendon midbody showing fascicular tendon (FT), peritendon (PT) and muscle fibres infiltrating the tendon structure. (E) Fascicular tendon fibrils shown with adjacent infiltrating muscle and adipocytes (A), also interfascicular matrix (IFM) and a vessel situated within the IFM (V). (F) Magnified view of fascicular tendon (FT) and peritendon (PT) from midbody showing adipocytes (A) and vessels (V) in the peritendon. G-I images are from the MTJ region. (G) Wide view of the MTJ where both fascicular tendon and muscle fibres are clearly visible, along with adipocytes. (H) Close view of the muscle-tendon interaction at the MTJ. (I) Close view of the fascicular tendon fibrils and inter-fascicular matrix (IFM). J-K Images from muscle. (J Wide view of muscle fibres (M) with stroma visible (S). (K) Higher magnification of myofibers. (L) Higher magnification of stroma showing artery (Ar), vein (V), peripheral nerve bundle (PNB) and adipocytes (Ad).

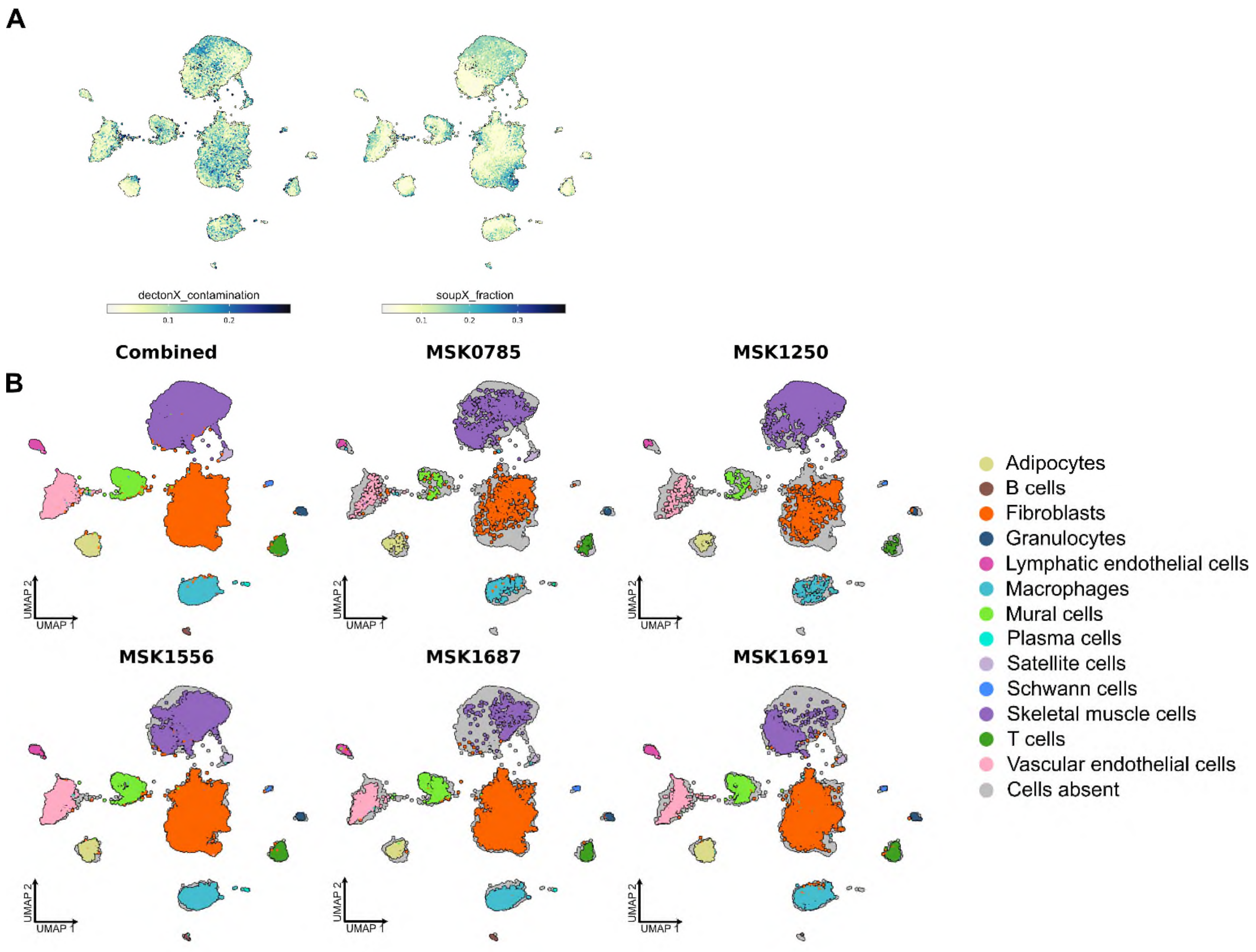


**Fig. S2. Quality control of snRNA-seq data.** (A) UMAP representation of cells from human Achilles tendon after filtering showing ambient RNA contamination measured by decontX (left) or soupX (right). (B) UMAP representation of cells from human Achilles tendon coloured by cell identity, with cells from each donor plotted in a separate panel.

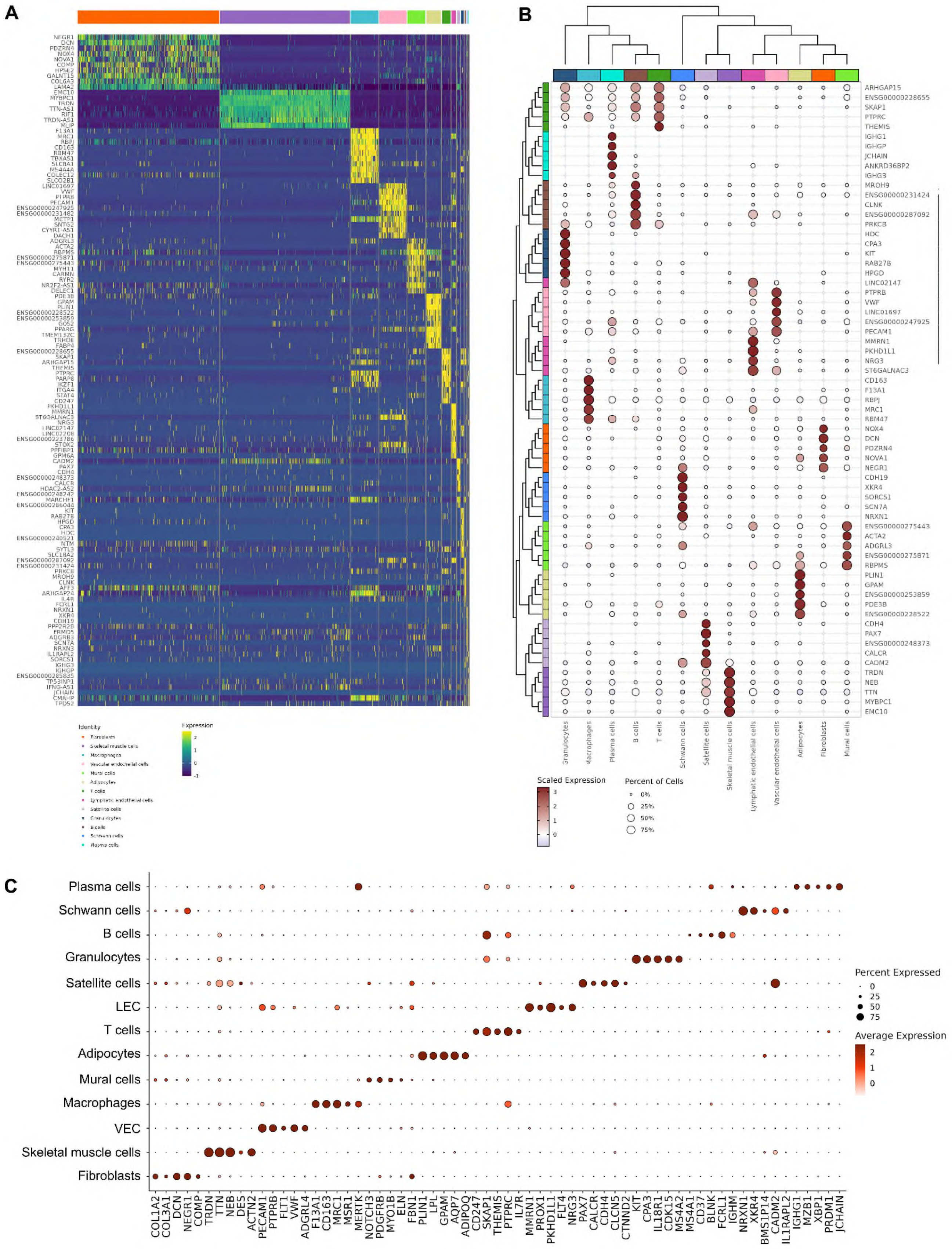
A
B
Scaled Expression
Percent of Cells
0%
25%
50%
75%
Granulocytes
Macrophages
Plasma cells
B cells
T cells
Schwann cells
Satellite cells
Skeletal muscle cells
Lymphatic endothelial cells
Vascular endothelial cells
Adipocytes
Fibroblasts
Mural cells
C
Plasma cells
Schwann cells
B cells
Granulocytes
Satellite cells
LEC
T cells
Adipocytes
Mural cells
Macrophages
VEC
Skeletal muscle cells
Fibroblasts
Percent Expressed
0
25
50
75
Average Expression
2
1
0
COL1A2
COL3A1
DCN
NEGR1
COMP
TRDN
TTN
NEB
DES
ACTN2
PECAM1
PTPRB
FLT1
VWF
ADGRL4
F13A1
CD163
MRC1
MSR1
MERTK
NOTCH3
PDGFRB
MYO1B
ELN
FBN1
PLIN1
LPL
GPAM
AQP7
ADIPOQ
CD247
SKAP1
THEMIS
PTPRC
IL7R
MMRN1
PROX1
PKHD1L1
FLT4
NRG3
PAX7
CALCR
CDH4
CLCN5
CTNND2
KIT
CPA3
IL18R1
CDK15
MS4A2
MS4A1
CD37
BLNK
FCRL1
IGHM
NRXN1
XKR4
BMS1P14
CADM2
IL1RAPL2
IGHG1
MZB1
XBP1
PRDM1
JCHAIN

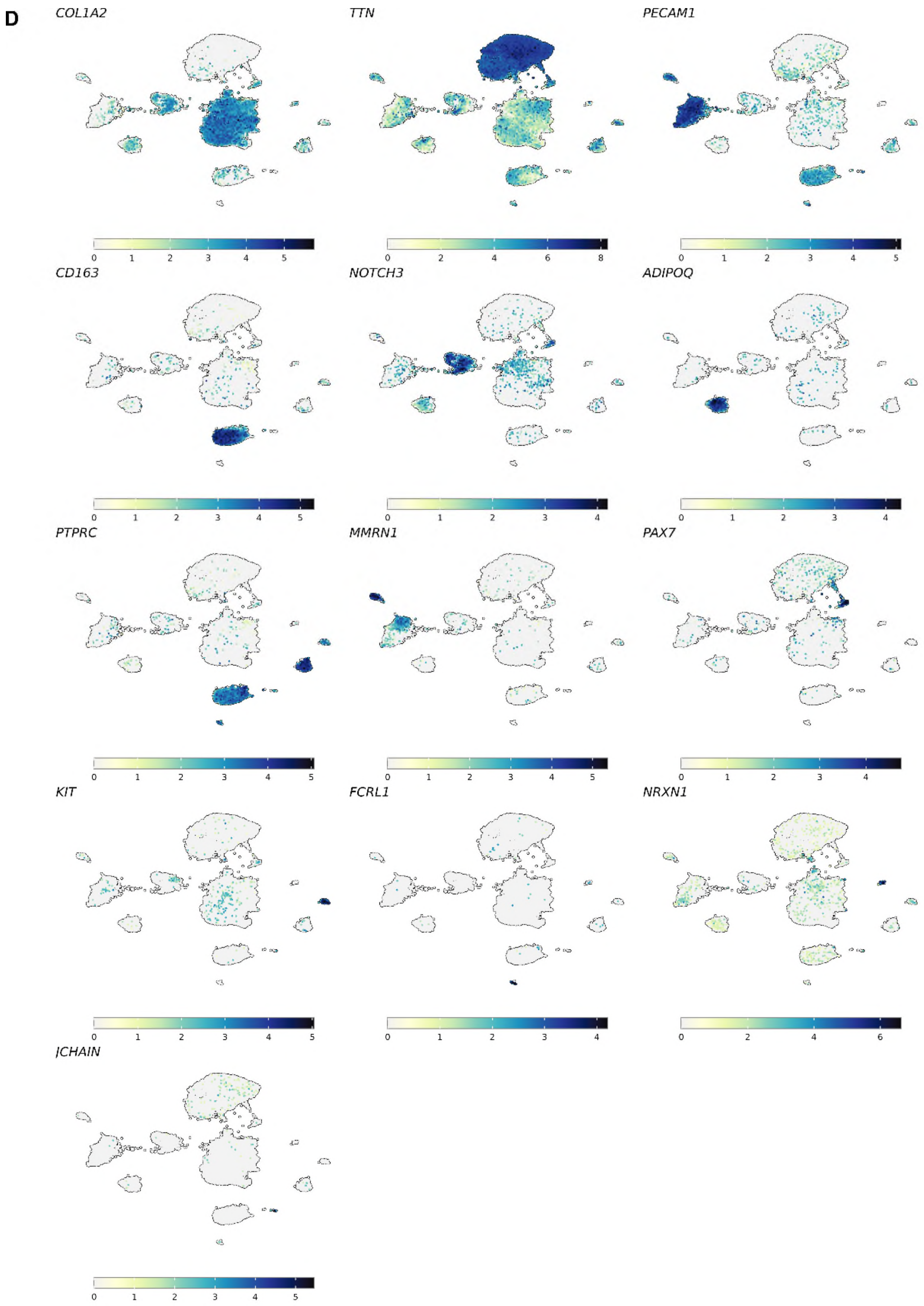


**Fig. S3. Annotation of snRNA-seq data.** (A) Heatmap showing scaled expression of top 10 differentially expressed genes for each broad cell type. (B) Clustered dotplot showing scaled expression of top 5 differentially expressed genes for each broad cell type. (C) Dotplot showing expression of canonical marker genes that were used to assign broad cell identities. (D) UMAP representations showing expression of canonical marker genes for each broad cell type.

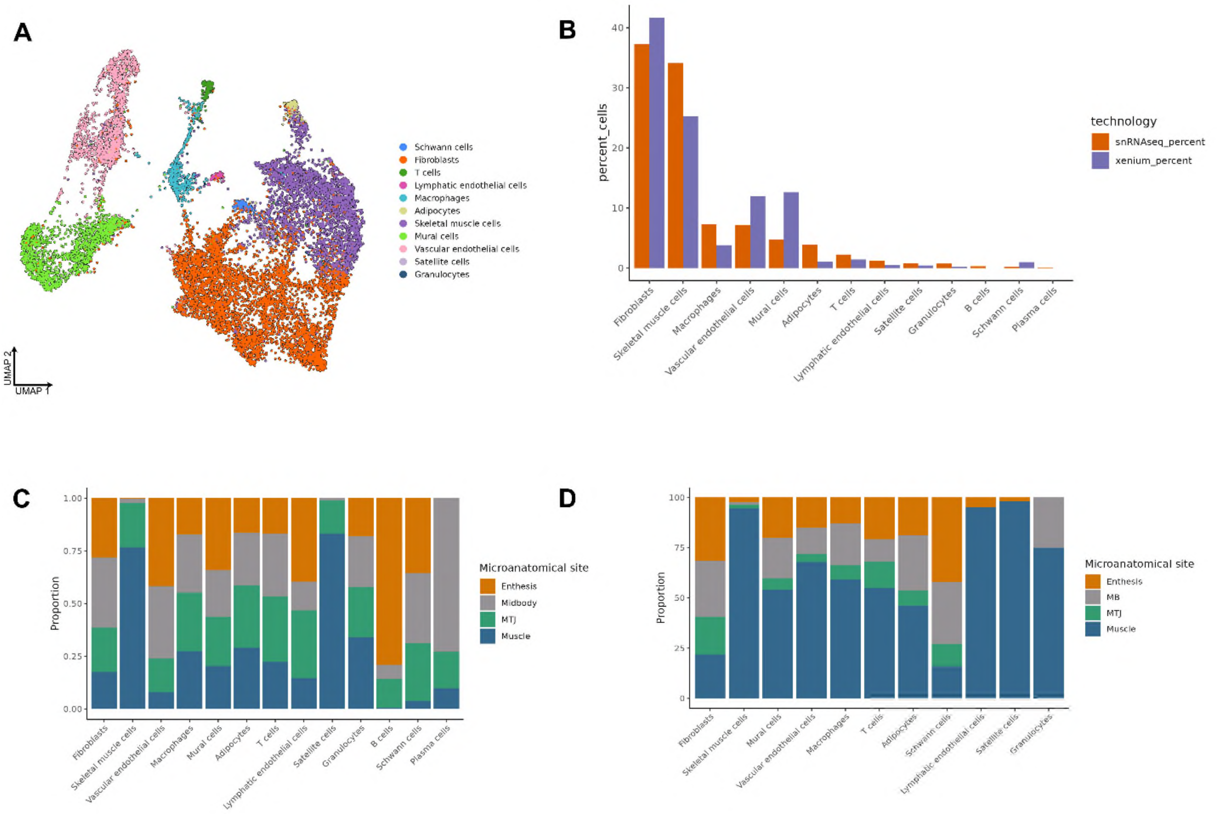


**Fig. S4. Comparison of cell types detected in snRNA-seq and Xenium data.** (A) UMAP representation of broad cell types in Xenium data, which were mapped from the snRNA-seq annotations. (B) Percentage of each cell type from snRNA-seq and Xenium data. (C) Proportion of each cell type present at each microanatomical site in snRNA-seq data. (D) Proportion of each cell type present at each microanatomical site in Xenium data.

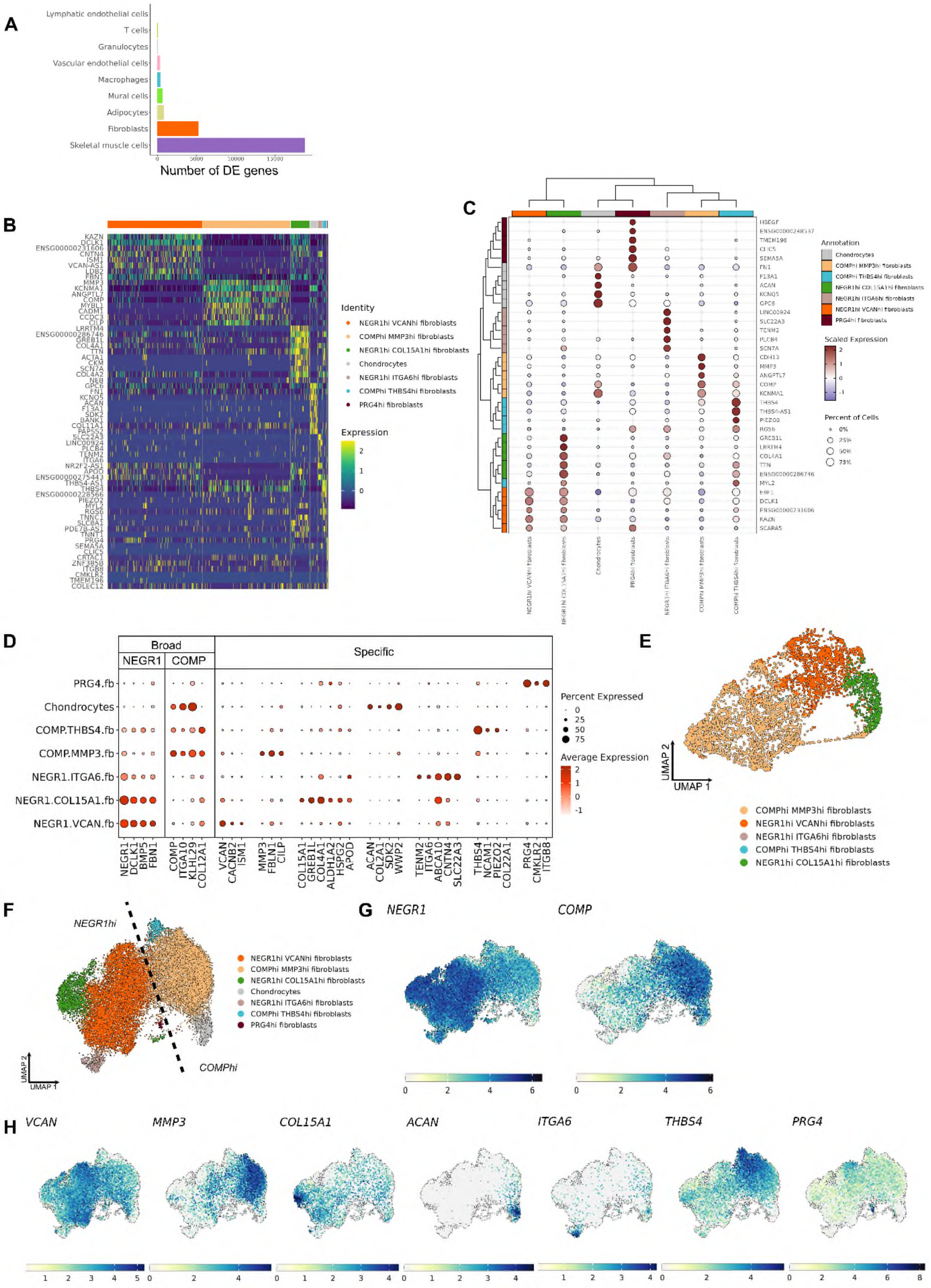

A
Lymphatic endothelial cells
T cells
Granulocytes
Vascular endothelial cells
Macrophages
Mural cells
Adipocytes
Fibroblasts
Skeletal muscle cells
Number of DE genes
B
Identity
NEGR1hi VCANhi fibroblasts
COMPhi MMP3hi fibroblasts
NEGR1hi COL15A1hi fibroblasts
Chondrocytes
NEGR1hi ITGA6hi fibroblasts
COMPhi THBS4hi fibroblasts
PRG4hi fibroblasts
Expression
C
Annotation
Chondrocytes
COMPhi MMP3hi fibroblasts
COMPhi THBS4hi fibroblasts
NEGR1hi COL15A1hi fibroblasts
NEGR1hi ITGA6hi fibroblasts
NEGR1hi VCANhi fibroblasts
PRG4hi fibroblasts
Scaled Expression
Percent of Cells
D
Broad
NEGR1
COMP
Specific
PRG4.fb
Chondrocytes
COMP.THBS4.fb
COMP.MMP3.fb
NEGR1.ITGA6.fb
NEGR1.COL15A1.fb
NEGR1.VCAN.fb
Percent Expressed
0
25
50
75
Average Expression
2
1
0
-1
NEGR1
DCLK1
BMP5
FBN1
COMP
ITGA10
KLHL29
COL12A1
VCAN
CACNB2
ISM1
MMP3
FBLN1
CILP
COL15A1
GREB1L
COL4A1
ALDH1A2
HSPG2
APOD
ACAN
COL2A1
SDK2
WWP2
TENM2
ITGA6
ABCA10
CNTN4
SLC22A3
THBS4
NCAM1
PIEZO2
COL22A1
PRG4
CMKLR2
ITGB8
E
UMAP 2
UMAP 1
COMPhi MMP3hi fibroblasts
NEGR1hi VCANhi fibroblasts
NEGR1hi ITGA6hi fibroblasts
COMPhi THBS4hi fibroblasts
NEGR1hi COL15A1hi fibroblasts
F
NEGR1hi
COMPhi
UMAP 2
UMAP 1
NEGR1hi VCANhi fibroblasts
COMPhi MMP3hi fibroblasts
NEGR1hi COL15A1hi fibroblasts
Chondrocytes
NEGR1hi ITGA6hi fibroblasts
COMPhi THBS4hi fibroblasts
PRG4hi fibroblasts
G
NEGR1
COMP
H
VCAN
MMP3
COL15A1
ACAN
ITGA6
THBS4
PRG4

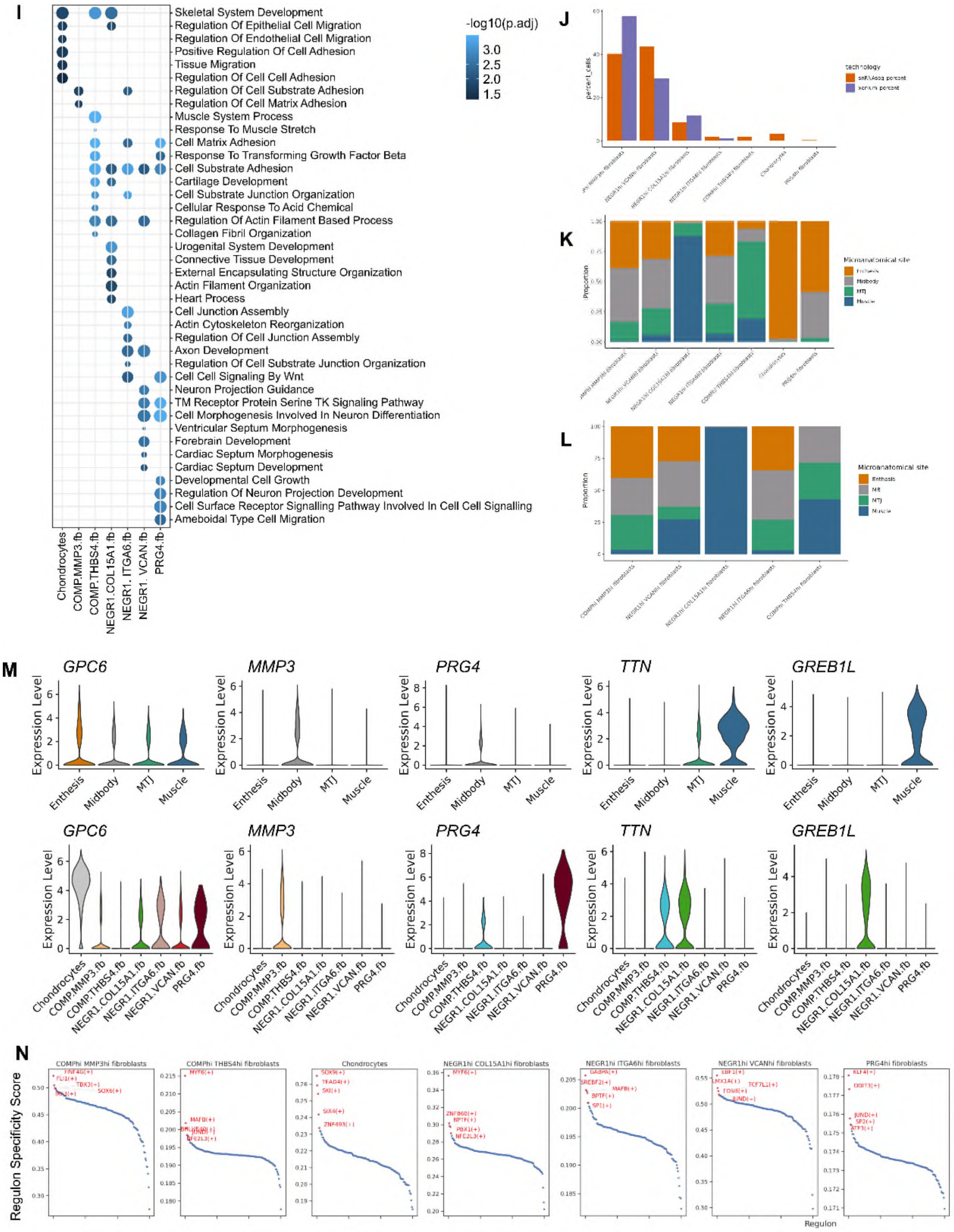
I
Skeletal System Development
Regulation Of Epithelial Cell Migration
Regulation Of Endothelial Cell Migration
Positive Regulation Of Cell Adhesion
Tissue Migration
Regulation Of Cell Cell Adhesion
Regulation Of Cell Substrate Adhesion
Regulation Of Cell Matrix Adhesion
Muscle System Process
Response To Muscle Stretch
Cell Matrix Adhesion
Response To Transforming Growth Factor Beta
Cell Substrate Adhesion
Cartilage Development
Cell Substrate Junction Organization
Cellular Response To Acid Chemical
Regulation Of Actin Filament Based Process
Collagen Fibril Organization
Urogenital System Development
Connective Tissue Development
External Encapsulating Structure Organization
Actin Filament Organization
Heart Process
Cell Junction Assembly
Actin Cytoskeleton Reorganization
Regulation Of Cell Junction Assembly
Axon Development
Regulation Of Cell Substrate Junction Organization
Cell Cell Signaling By Wnt
Neuron Projection Guidance
TM Receptor Protein Serine TK Signaling Pathway
Cell Morphogenesis Involved In Neuron Differentiation
Ventricular Septum Morphogenesis
Forebrain Development
Cardiac Septum Morphogenesis
Cardiac Septum Development
Developmental Cell Growth
Regulation Of Neuron Projection Development
Cell Surface Receptor Signalling Pathway Involved In Cell Cell Signalling
Ameboidal Type Cell Migration
Chondrocytes
COMP.MMP3.fb
COMP.THBS4.fb
NEGR1.COL15A1.fb
NEGR1. ITGA6.fb
NEGR1. VCAN.fb
PRG4.fb
-log10(p.adj)
3.0
2.5
2.0
1.5
J
K
L
Microanatomical site
Proportion
M
GPC6
MMP3
PRG4
TTN
GREB1L
Expression Level
Enthesis
Midbody
MTJ
Muscle
Chondrocytes
COMP.MMP3.fb
COMP.THBS4.fb
NEGR1.COL15A1.fb
NEGR1.ITGA6.fb
NEGR1.VCAN.fb
PRG4.fb
N
Regulon Specificity Score
Regulon

**Fig. S5. Fine annotation of fibroblast subset.** (A) Number of differentially expressed genes between midbody/MTJ/muscle compared to enthesis in each broad cell type. (B) Heatmap showing scaled expression of top 10 differentially expressed genes for each finely annotated fibroblast cell type in snRNA-seq data. (C) Clustered dotplot showing scaled expression of top 5 differentially expressed genes for each finely annotated fibroblast cell type in snRNA-seq data. (D) Dotplot showing expression of broad and specific marker genes used to assign finely annotated cell identities in the fibroblast subset in snRNA-seq data. (E) UMAP representation of fibroblast cell types in Xenium data. (F) UMAP representations showing fibroblast subtypes demonstrating the two major fibroblast groups expressing NEGR1 and COMP. (G) UMAP representation showing expression of broad markers *NEGR1*and *COMP* in the fibroblast subset (H) UMAP representations showing expression of marker genes for each fibroblast subtype. (I) Dotplot showing GO biological pathways enriched in each fibroblast subtype. (J) Percentage of each fibroblast cell type from snRNA-seq and Xenium data. (K) Average proportion of each fibroblast cell type present at each microanatomical site in snRNA-seq data. (L) Proportion of each fibroblast cell type present at each microanatomical site in Xenium data. (M) Violin plots showing expression of selected genes across microanatomy (upper panel) and in fibroblast subtypes (lower panel). (N) Results of SCENIC analysis showing regulon specificity scores in each fibroblast subtypes, highlighting the top 5 most specific regulons.

Abbreviations: PRG4.fb - PRG4hi fibroblasts; COMP.THBS4.fb – COMPhi THBS4hi fibroblasts; NEGR1.ITGA6.fb – NEGR1hi ITGA6hi fibroblasts; NEGR1.COL15A1.fb – NEGR1hi COL15A1 fibroblasts; COMP.MMP3.fb – COMPhi MMP3hi fibroblasts; NEGR1.VCAN.fib - NEGR1hi VCANhi fibroblasts.

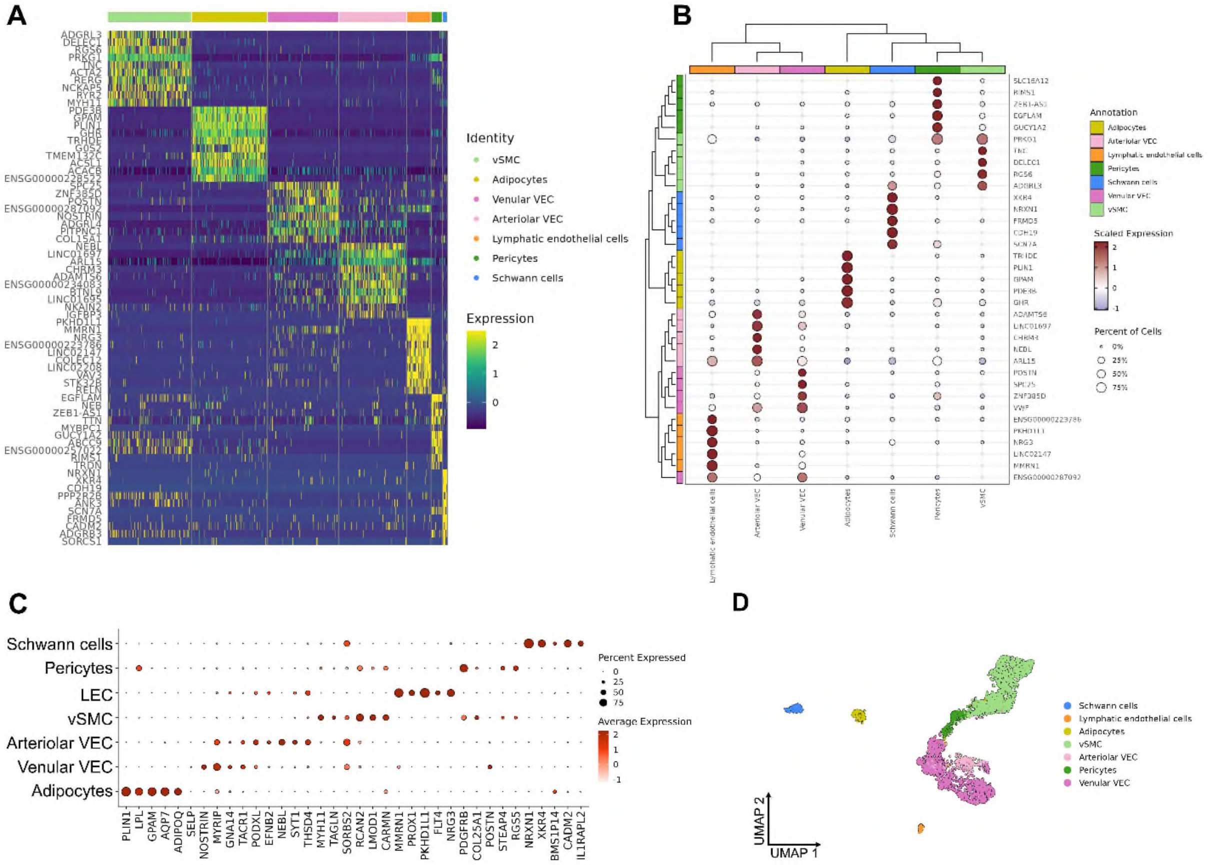

**Fig. S6. Fine annotation of stromal cell types.** (A) Heatmap showing scaled expression of top 10 differentially expressed genes for each finely annotated skeletal muscle cell type in snRNA-seq data. (B) Clustered dotplot showing scaled expression of top 5 differentially expressed genes for each finely annotated stromal cell type in snRNA-seq data. (C) Dotplot showing expression of canonical marker genes used to assign finely annotated cell identities in each stromal cell subset in snRNA-seq data. (D) UMAP representation of stromal cell types in Xenium data.

A

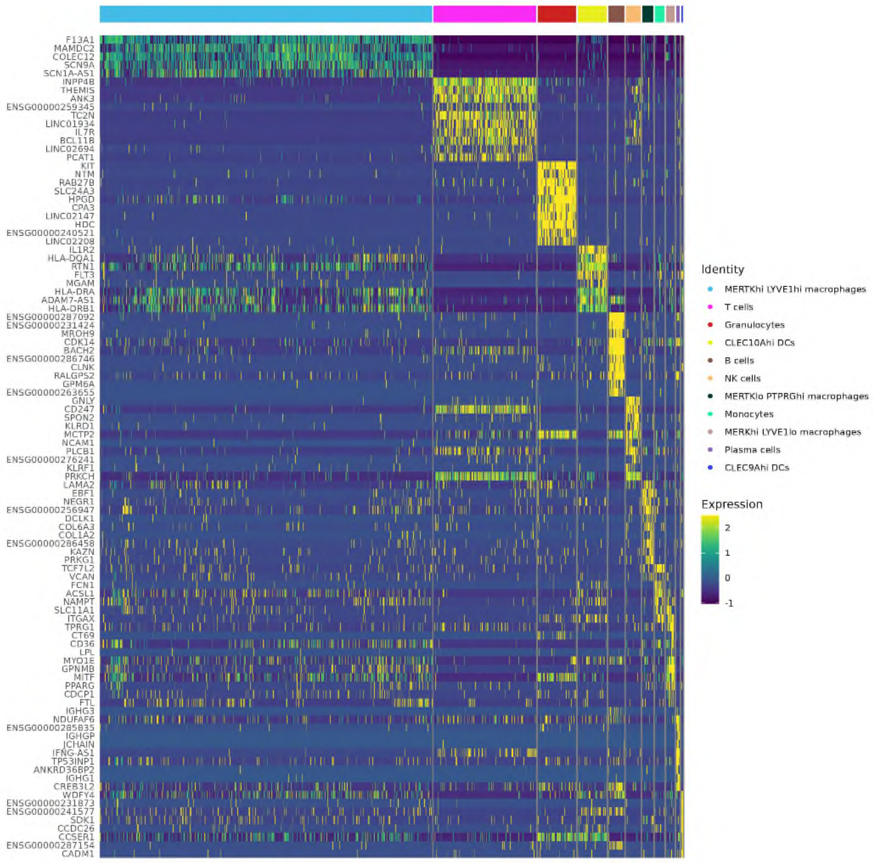
F13A1
MAMDC2
COLEC12
SCN9A
SCN1A-AS1
INPP4B
THEMIS
ANK3
ENSG00000259345
TC2N
LINC01934
IL7R
BCL11B
LINC02694
PCAT1
KIT
NTM
RAB27B
SLC24A3
HPGD
CPA3
LINC02147
HDC
ENSG00000240521
LINC02208
IL1R2
HLA-DQA1
RTN1
FLT3
MGAM
HLA-DRA
ADAM7-AS1
HLA-DRB1
ENSG00000287092
ENSG00000231424
MROH9
CDK14
BACH2
ENSG00000286746
CLNK
RALGPS2
GPM6A
ENSG00000263655
GNLY
CD247
SPON2
KLRD1
MCTP2
NCAM1
PLCB1
ENSG00000276241
KLRF1
PRKCH
LAMA2
EBF1
NEGR1
ENSG00000256947
DCLK1
COL6A3
COL1A2
ENSG00000286458
KAZN
PRKG1
TCF7L2
VCAN
FCN1
ACSL1
NAMPT
SLC11A1
ITGAX
TPRG1
CT69
CD36
LPL
MYO1E
GPNMB
MITF
PPARG
CDCP1
FTL
IGHG3
NDUFAF6
ENSG00000285835
IGHGP
JCHAIN
IFNG-AS1
TP53INP1
ANKRD36BP2
IGHG1
CREB3L2
WDFY4
ENSG00000231873
ENSG00000241577
SDK1
CCDC26
CCSER1
ENSG00000287154
CADM1
Identity
MERTKhi LYVE1hi macrophages
T cells
Granulocytes
CLEC10Ahi DCs
B cells
NK cells
MERTKlo PTPRGhi macrophages
Monocytes
MERKhi LYVE1lo macrophages
Plasma cells
CLEC9Ahi DCs
Expression
2
1
0
-1

B

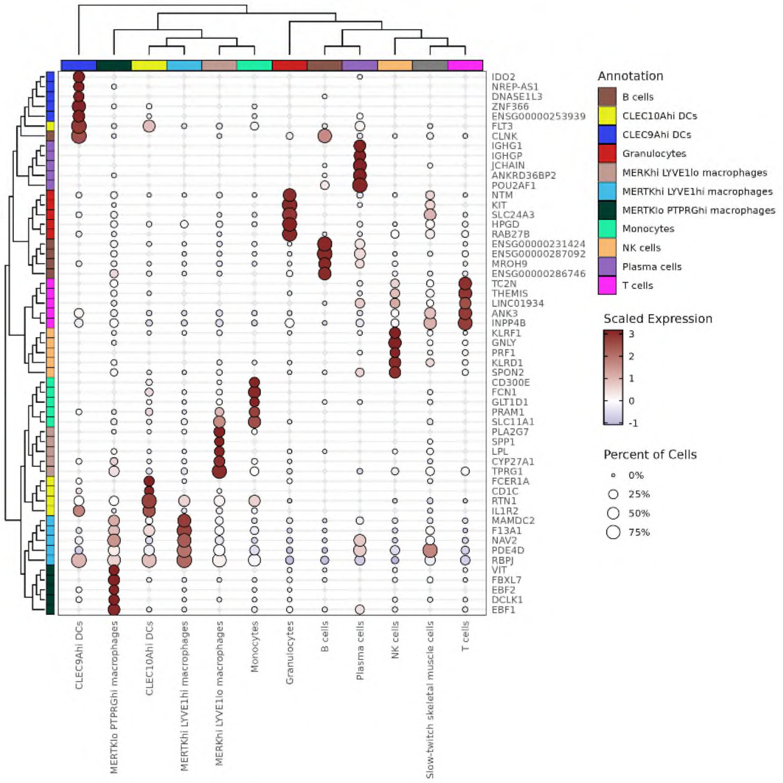
IDO2
NREP-AS1
DNASE1L3
ZNF366
ENSG00000253939
FLT3
CLNK
IGHG1
IGHGP
JCHAIN
ANKRD36BP2
POU2AF1
NTM
KIT
SLC24A3
HPGD
RAB27B
ENSG00000231424
ENSG00000287092
MROH9
ENSG00000286746
TC2N
THEMIS
LINC01934
ANK3
INPP4B
KLRF1
GNLY
PRF1
KLRD1
SPON2
CD300E
FCN1
GLT1D1
PRAM1
SLC11A1
PLA2G7
SPP1
LPL
CYP27A1
TPRG1
FCER1A
CD1C
RTN1
IL1R2
MAMDC2
F13A1
NAV2
PDE4D
RBPJ
VIT
FBXL7
EBF2
DCLK1
EBF1
CLEC9Ahi DCs
MERTKlo PTPRGhi macrophages
CLEC10Ahi DCs
MERTKhi LYVE1hi macrophages
MERKhi LYVE1lo macrophages
Monocytes
Granulocytes
B cells
Plasma cells
NK cells
Slow-twitch skeletal muscle cells
T cells
Annotation
B cells
CLEC10Ahi DCs
CLEC9Ahi DCs
Granulocytes
MERKhi LYVE1lo macrophages
MERTKhi LYVE1hi macrophages
MERTKlo PTPRGhi macrophages
Monocytes
NK cells
Plasma cells
T cells
Scaled Expression
3
2
1
0
-1
Percent of Cells
0%
25%
50%
75%

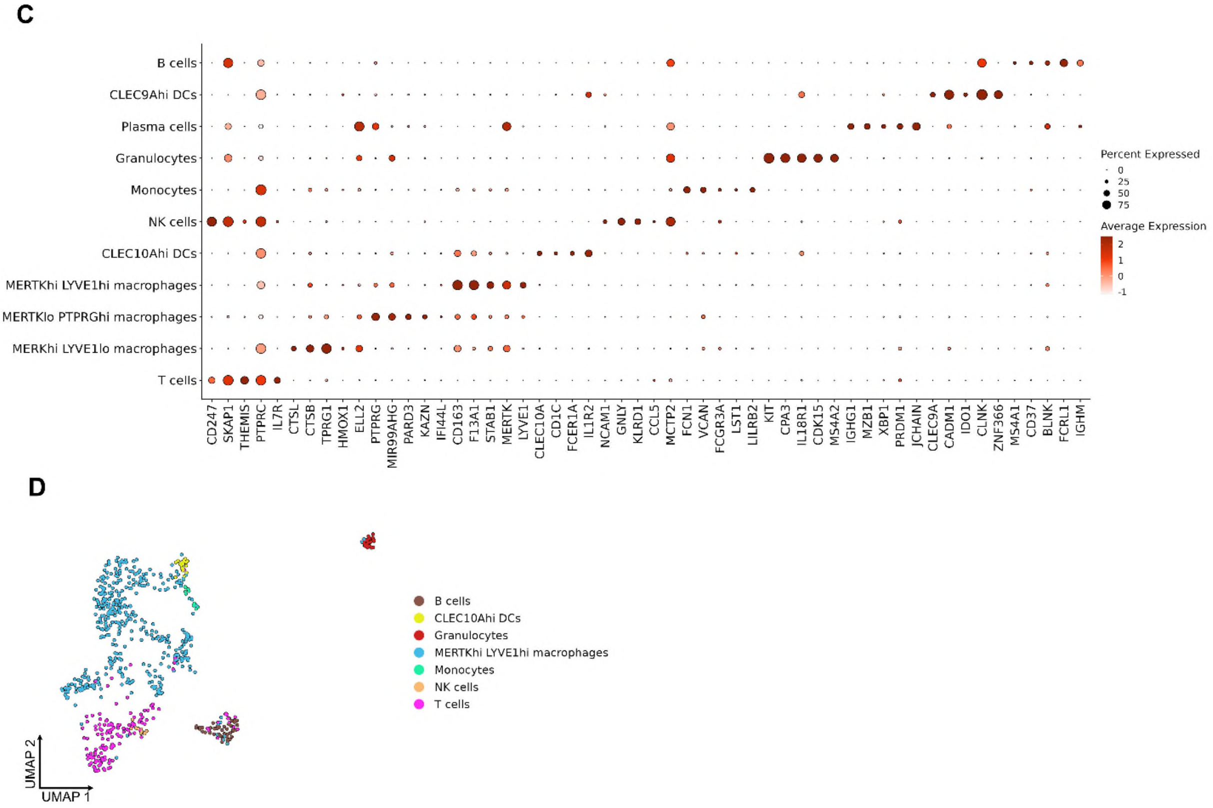


**Fig. S7. Fine annotation of immune cells.** (A) Heatmap showing scaled expression of top 10 differentially expressed genes for each finely annotated immune cell type in snRNA-seq data. (B) Clustered dotplot showing scaled expression of top 5 differentially expressed genes for each finely annotated immune cell type in snRNA-seq data. (C) Dotplot showing expression of canonical marker genes used to assign finely annotated cell identities in each immune cell subset in snRNA-seq data (D) UMAP representation of immune cell types in Xenium data.

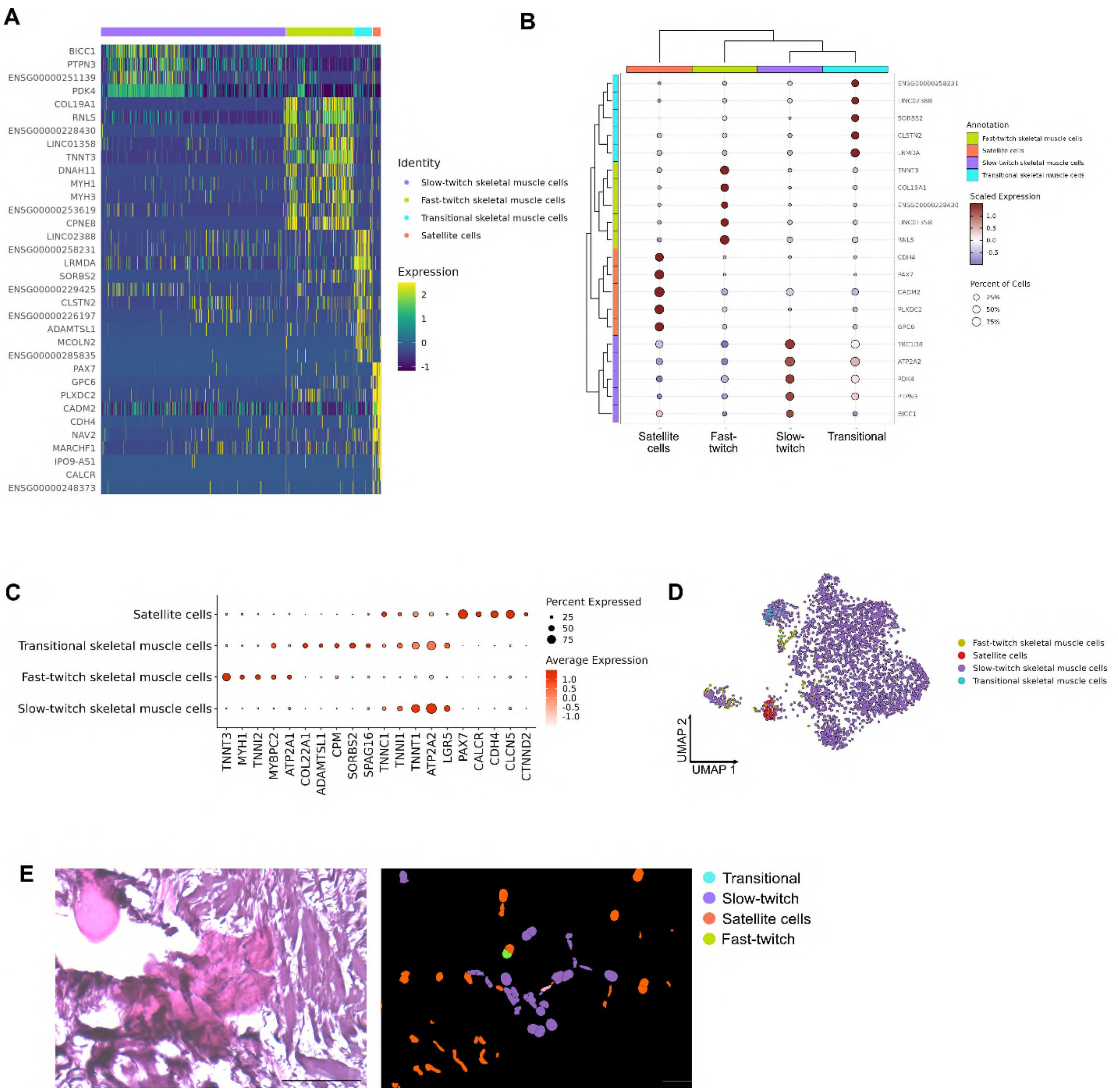


**Fig. S8. Fine annotation of muscle cells.** (A) Heatmap showing scaled expression of top 10 differentially expressed genes for each finely annotated skeletal muscle cell type in snRNA-seq data. (B) Clustered dotplot showing scaled expression of top 5 differentially expressed genes for each finely annotated skeletal muscle cell type in snRNA-seq data. (C) Dotplot showing expression of canonical marker genes used to assign finely annotated cell identities in each skeletal muscle cell subset in snRNA-seq data (D) UMAP representation of skeletal muscle cell types in Xenium data. (E) H&E image (left) and spatial transcriptomics plot (right) of the Achilles midbody, showing muscle fibres and cells present in the fascicular tendon matrix. Scale bar 100 µm.

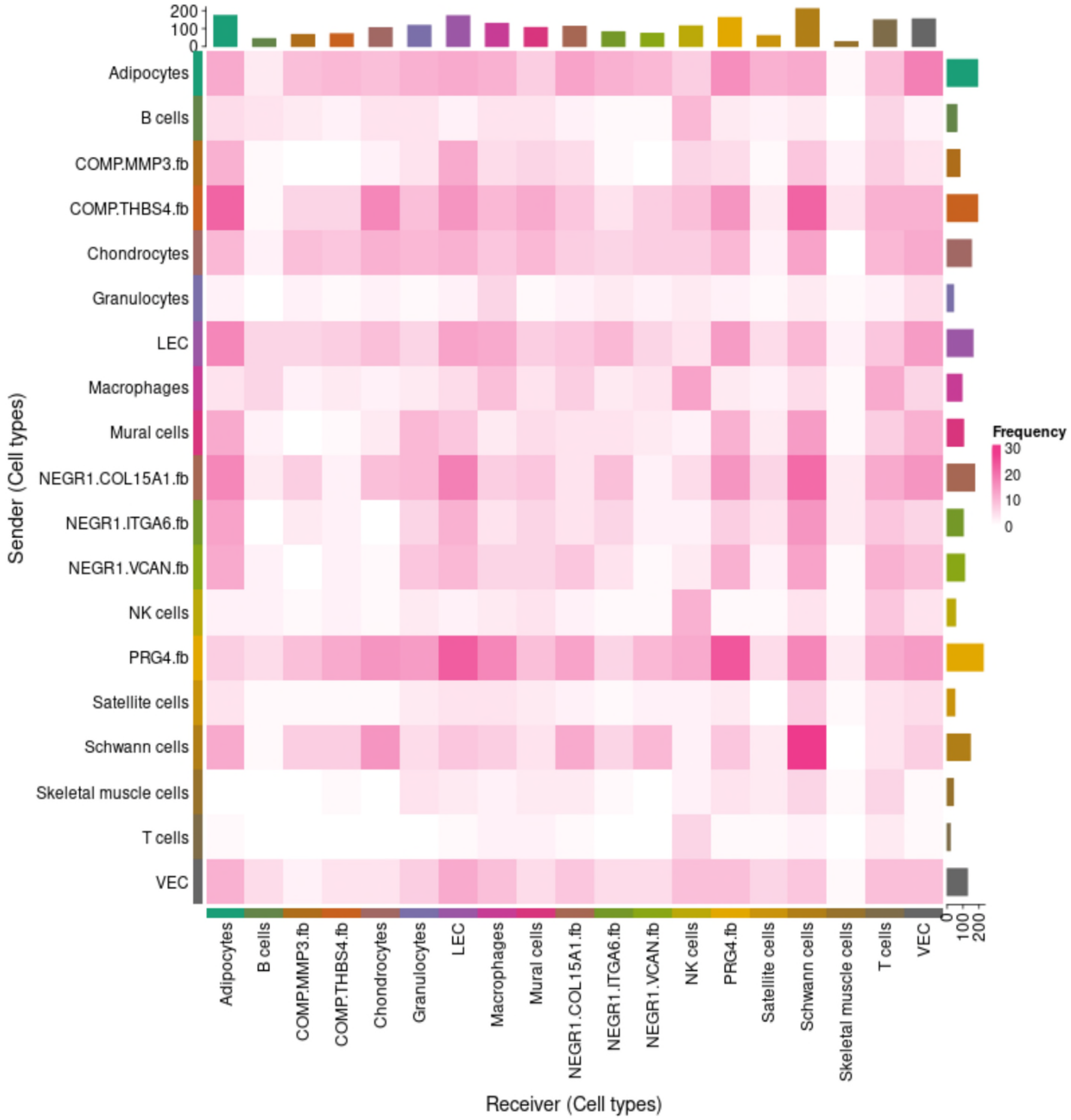


**Fig. S9. Heatmap showing frequency of cell-cell interactions in Achilles tendon**. Abbreviations as Figure 4.

**Data S1. (separate file)**

Differentially expressed genes for each broadly annotated cell type compared to all other cell types identified using Seurat FindMarkers(). P-values: significance using Wilcoxon Rank Sum test; avg_logFC: log fold-chage of the average expression between the two groups. Positive values indicate that the gene is more highly expressed in the first group; pct.1: The percentage of cells where the gene is detected in the first group; pct.2: The percentage of cells where the gene is detected in the second group; p_val_adj: Adjusted p-value, based on bonferroni correction using all genes in the dataset; cluster: cell type; gene: human readable gene ID or EnsemblID.

**Data S2. (separate file)**
Differentially expressed genes for each finely annotated cell type compared to all other cell types identified using Seurat FindMarkers(). Columns as for Data S1.

**Data S3. (separate file).**
Gene names, Ensemble IDs and number of probes used in the Xenium Add-On Custom Probe Panel.
